# Regional specialization manifests in the reliability of neural population codes

**DOI:** 10.1101/2024.01.25.576941

**Authors:** Jennifer A. Guidera, Daniel P. Gramling, Alison E. Comrie, Abhilasha Joshi, Eric L. Denovellis, Kyu Hyun Lee, Jenny Zhou, Paige Thompson, Jose Hernandez, Allison Yorita, Razi Haque, Christoph Kirst, Loren M. Frank

## Abstract

The brain has the remarkable ability to learn and guide the performance of complex tasks. Decades of lesion studies suggest that different brain regions perform specialized functions in support of complex behaviors^1–3^. Yet recent large-scale studies of neural activity reveal similar patterns of activity and encoding distributed widely throughout the brain^4–6^. How these distributed patterns of activity and encoding are compatible with regional specialization of brain function remains unclear. Two frontal brain regions, the dorsal medial prefrontal cortex (dmPFC) and orbitofrontal cortex (OFC), are a paradigm of this conundrum. In the setting complex behaviors, the dmPFC is necessary for choosing optimal actions^2,7,8^, whereas the OFC is necessary for waiting for^3,9^ and learning from^2,7,9–12^ the outcomes of those actions. Yet both dmPFC and OFC encode both choice- and outcome-related quantities^13–20^. Here we show that while ensembles of neurons in the dmPFC and OFC of rats encode similar elements of a cognitive task with similar patterns of activity, the two regions differ in when that coding is consistent across trials (“reliable”). In line with the known critical functions of each region, dmPFC activity is more reliable when animals are making choices and less reliable preceding outcomes, whereas OFC activity shows the opposite pattern. Our findings identify the dynamic reliability of neural population codes as a mechanism whereby different brain regions may support distinct cognitive functions despite exhibiting similar patterns of activity and encoding similar quantities.

## RESULTS

We simultaneously monitored the activity of large neuronal ensembles in the dmPFC and OFC of rats as they performed a cognitive task that involves choosing among options, which requires the dmPFC^2,8,21^, and waiting for^3,9^ and adapting behavior based on^2,7,9–12^ the outcomes of choices, which requires the OFC (Fig. 1a–e). On each trial, rats chose among reward wells in a maze and expressed their choice by traversing the path to their chosen well (“path traversal”) (Fig. 1a). Following arrival to the well, rats waited a set amount of time (two seconds) before learning the outcome of their choice (“delay period”), either a milk reward or its omission per an alternation rule (Fig. 1a,b). Rats performed the task across different environments and with different spatial arrangements of reward wells (Fig. 1c, Extended Data Fig. 1a). Here we focus on activity recorded after rats had learned to perform the task across environments and sets of reward wells (Extended Data Fig. 1a,b, Supplementary Video 1). Simultaneous recordings from the dorsal hippocampus (HPc) in a subset of subjects provided a point of comparison to an area known to exhibit reliable coding in rats performing cognitive tasks^22,23^ (Fig. 1d).

**Fig. 1.**
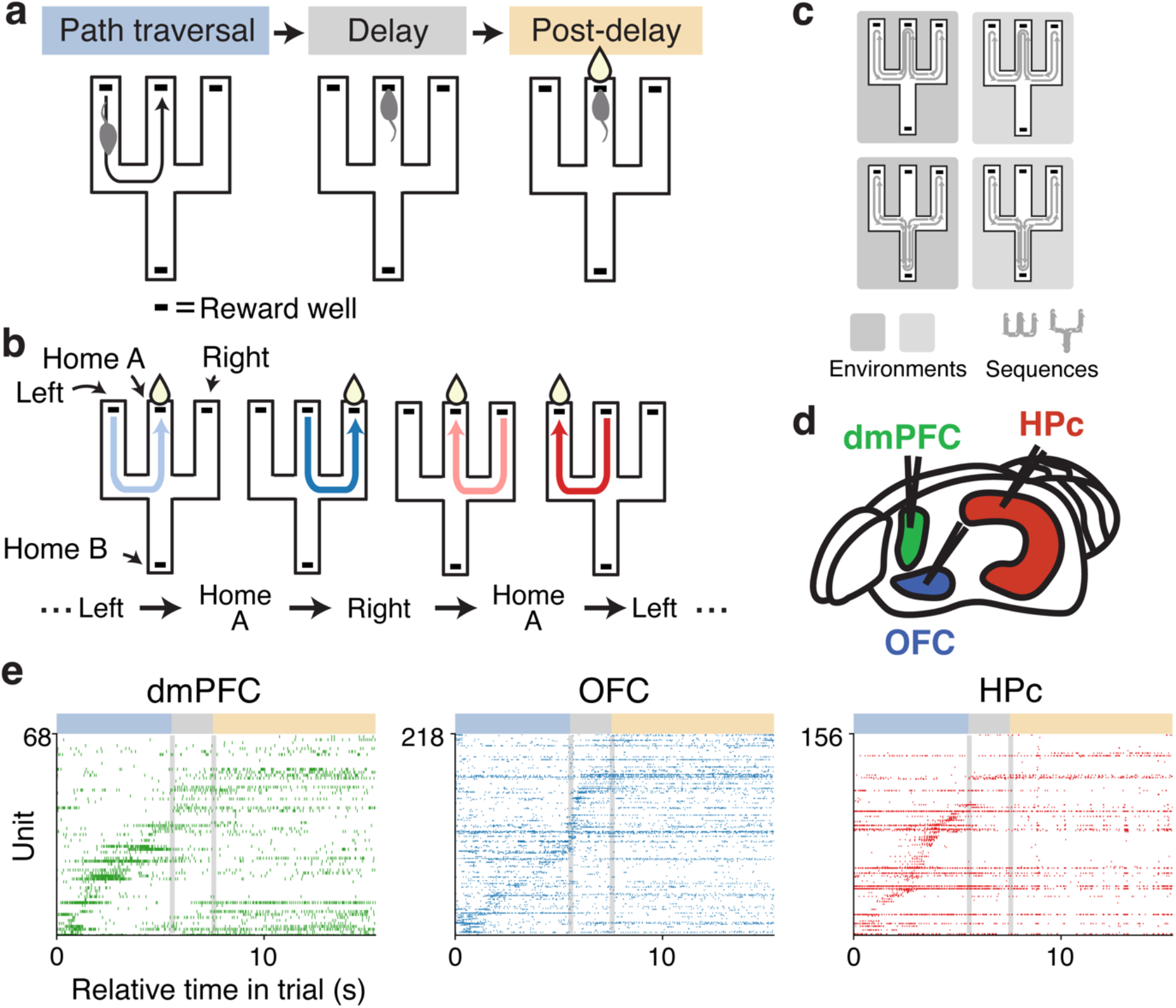
Multisite recordings in rats performing a cognitive task. **a**, Trial structure. On each trial, rats navigate to a chosen well (“path traversal”) where they wait two seconds before learning the outcome of their choice (“delay”). Rats self-initiate a new trial by leaving the well. **b**, Task rule. Rats receive a milk reward for making alternating visits to the left and right wells from the designated home well. A sequence of correct well visits is illustrated with home A as the designated home well. **c**, Rats performed the alternation task across different environments and with different sets of reward wells (sequences). **d**, Recording sites. **e**, Rasters of simultaneously recorded dmPFC, OFC, and HPc single units during a single trial in an example rat. Colored rectangles and vertical lines mark the task phases in **a**.

### Embeddings suggest dmPFC and OFC express a common code for task abstractions with distinct patterns of reliability

While previous work identified broadly similar patterns of task-related activity in single dmPFC and OFC neurons^4,13–15,24^, whether the population-level organization of neural activity is similar or different across these regions was unknown. We first addressed that question by visualizing the activity of hundreds of simultaneously recorded dmPFC and OFC neurons as rats performed the task. A standard linear dimensionality reduction approach (principal component analysis)^25,26^ failed to yield a low-dimensional subspace that captured a substantial fraction of the variance in the firing rates across simultaneously recorded neurons in each brain region (Extended Data Fig. 2b). We therefore turned to a method that can capture low-dimensional, non-linear structure in data, uniform manifold approximation and projection (UMAP)^27^.

Applying a 3-dimensional UMAP to 100-ms firing rate vectors from neurons recorded in each brain region across entire behavioral sessions suggested an intriguing possibility: that dmPFC and OFC encode similar task-related quantities but express this common code reliably at distinct task phases associated with either choice (dmPFC) or outcomes (OFC) (Fig. 2, Supplementary Video 2). Beginning with the common code, embedded activity in both dmPFC and OFC was structured around the act of goal-seeking, followed by the specific settings in which goal-seeking took place. In both regions, the greatest variation in embedded activity mapped the abstract progression towards goals (potential reward), both as rats traversed the path and then waited to learn the outcome of their choice during the delay period at the well (Fig. 2a,b). Within this global progression, embedded activity diverged along paths requiring distinct turn sequences (Fig. 2c), thus mapping the actions required to reach reward. A comparison to HPc embedded activity underscored the relative similarity of the dmPFC and OFC codes. In contrast to an organization around the abstract progression towards goals, HPc embedded activity was primarily organized around the specific progressions along individual paths and in the delay period at individual reward wells (Fig. 2a–c), consistent with the well-established spatial and directional selectivity of HPc neurons^23,28^.

**Fig. 2.**
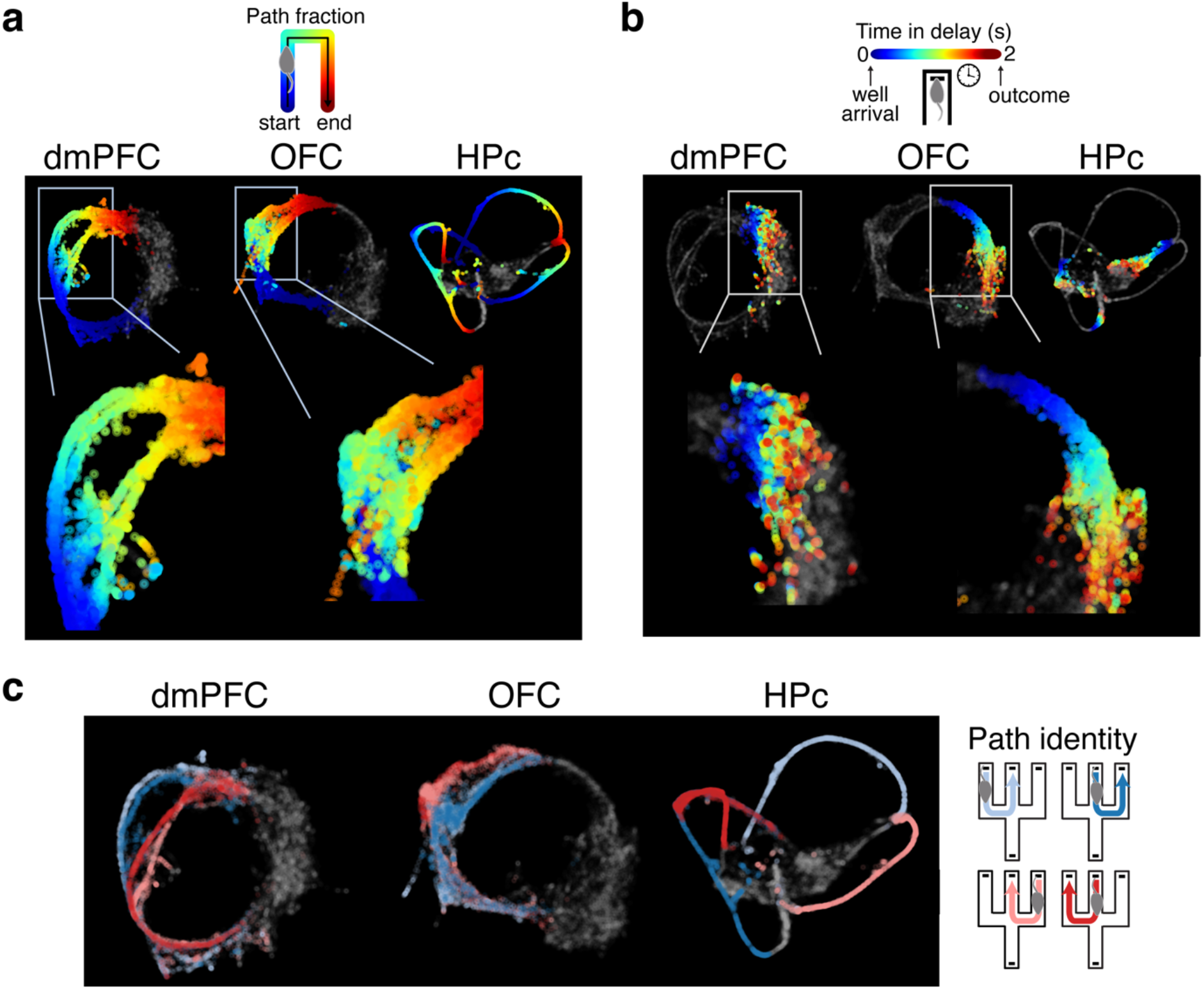
Population activity embeddings suggest dmPFC and OFC express a common code for the task with distinct patterns of reliability. **a–c**, UMAP embeddings of dmPFC (left), OFC (middle), and HPc (right) 100-ms firing rate vectors during an example session from one rat, colored by (**a**) the fractional distance along paths during path traversals, (**b**) relative time in the delay period, or (**c**) path identity during path traversals. Insets expand embeddings in the regions marked by rectangles to highlight differences between dmPFC and OFC.

The embeddings further suggested that dmPFC and OFC specialize by reliably expressing this common code at either choice-related (dmPFC) or outcome-related (OFC) phases of the task. Consistent activity across trials (“reliability”) manifests as more concentrated regions of the embedding. In the dmPFC, embedded activity evolved in a highly consistent manner as rats actively expressed choices by traversing paths to chosen wells (Fig. 2a). During the same period, OFC activity was much more diffuse, particularly towards the middle of paths (Fig. 2a). Conversely, as animals approached reward wells where outcomes are revealed and then waited to learn the outcomes of their choices, embedded activity precisely tracked task progression in the OFC (Fig. 2b). The same period was associated with much more diffuse activity patterns in the dmPFC (Fig. 2b).

These same overall patterns of task coding and the same differences in reliability across brain regions were present across animals, environments, and different sets of reward wells (Extended Data Fig. 3a–f). Altogether, the embeddings thus led us to hypothesize that dmPFC and OFC encode similar task-related quantities with similar average patterns of activity but express this common code reliably at distinct task phases to support distinct computations. We next set out to test these hypotheses quantitatively with both single cell and population analyses.

### A similar single unit code is reliable at distinct task phases across dmPFC and OFC

The embeddings suggested that dmPFC and OFC share a common code for the task that primarily maps the abstract progression towards goals, and secondarily maps the sensorimotor specifics that distinguish those progressions. An examination of the activity of single units (putative single neurons) in dmPFC and OFC revealed firing patterns consistent with this possibility. Dorsal mPFC and OFC single units tended to show the most prominent firing rate modulations over the course of a trial, and modulations across different paths or at different reward wells during the delay period tended to be similar (Fig. 3a,b), a result consistent with previous work^17,18,29–32^. By contrast, HPc single units largely showed distinct firing rate modulations across these conditions, consistent with the well-established location and heading direction selectivity of HPc neurons (Fig. 3c)^28,30^.

**Fig. 3.**
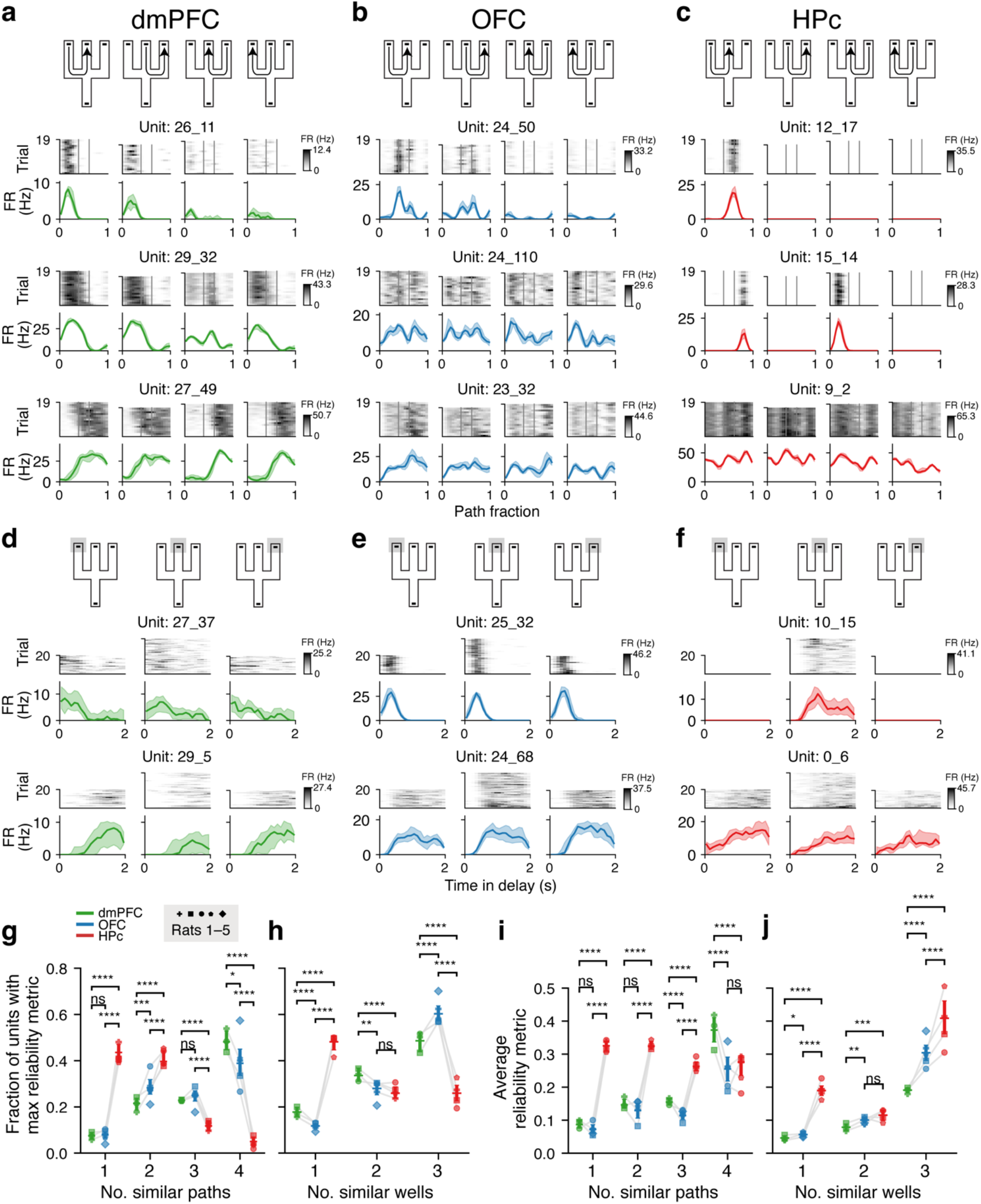
A common code for the task is reliable at different task phases in dmPFC versus OFC single neurons. **a–f**, Firing rates of example dmPFC (**a,d**), OFC (**b,e**), and HPc (**c,f**) single units from a single subject during path traversals (**a–c**) or the delay period at wells (**d–f**). Each panel shows single-trial firing rates (top) and corresponding mean and interquartile range (bottom) for an example single unit along paths (**a–c**) or during the delay period wells (**d–f**). **g**, Fraction of single units in each brain region that were best described as representing progression along individual paths or generalized across two, three, or four paths. Assignments were based on whether the summed reliability metric during path traversals was greatest for one, two, three, or four paths. In g–j: Horizontal lines show the mean across single units from 105 sessions across rats (dmPFC, *n* = 3 rats; OFC, *n* = 5 rats; HPc, *n* = 4 rats), and bars show 95% confidence intervals (CIs). Markers show the mean across sessions and single units for individual rats. Gray lines connect means for individual rats. * p < 0.05, ** p < 0.01, *** p < 0.001, **** p < 0.0001; ns, not significant (p ≥ 0.05). **h**, Fraction of single units in each brain region that were best described as representing progression in the delay period at individual wells or generalized across two or three wells. Assignments are based on whether the summed reliability metric during the delay period was greatest for one, two, or three wells. **i**, Average reliability metric across single units for which the summed reliability metric was greatest for one, two, three, or four paths during path traversals. **j**, Average reliability metric across single units for which the summed reliability metric was greatest for one, two, or three wells during the delay period.

To quantify the reliability of these patterns, we developed a metric (Extended Data Fig. 4a–d) that provides information about the extent to which units reliably encode progression towards goals uniquely within specific spatial and sensorimotor contexts versus similarly across contexts. For path traversals this yields four values, corresponding to the reliability of path progression representations shared across one, two, three, or all four paths (Extended Data Fig. 4c). The absolute magnitude of each value reflects the overall reliability of the activity across that number of paths, with higher values indicating greater predictability of single trial activity.

The values of these metrics from a single behavioral session revealed evidence for both a common code across dmPFC and OFC at the level of single neurons and differences between the regions in when that code was reliably expressed (Extended Data Fig. 4k–n). Single units in both dmPFC and OFC frequently showed reliable activity patterns that were similar across two, three, or four paths, with the greatest number of single units showing reliable firing generalized across all four paths (Extended Data Fig. 4k). At the same time, the magnitude of the values was notably higher in dmPFC than in OFC, reflecting more reliable task progression representations in dmPFC as rats traversed paths (Extended Data Fig. 4k). During the delay period, single dmPFC and OFC units likewise showed increasingly reliable firing rate patterns across increasing numbers of wells, but values were higher in OFC than in dmPFC, reflecting more reliable task progression representations in the OFC as rats waited for outcomes at wells (Extended Data Fig. 4l). This coding scheme was distinct from that in the HPc, where units most often showed reliable activity patterns unique to one or two paths during path traversals (Extended Data Fig. 4k) and one well during the delay period (Extended Data Fig. 4l).

The same results were seen across the full datasets from all five animals (Fig. 3g–j). Both dmPFC and OFC single units showed more reliable task progression coding across greater numbers of paths, whereas the opposite was seen in the HPc (Fig. 3g). Moreover, dmPFC reliability values were higher than OFC values for the many neurons that showed similar firing across three or four paths (Fig. 3i). And both dmPFC and OFC had higher reliability values for two and three wells across the delay period (Fig. 3h), but OFC values were higher than those for dmPFC (Fig. 3j).

### A similar population-level code is reliable at distinct task phases across dmPFC and OFC

The consistent patterns of results seen in the single cell analyses motivated a parallel set of analyses applied to the population of simultaneously recorded units in each brain region. Our first goal was to quantify, at the population level, the extent to which task progression coding during choice-related (path traversals) and outcome-related (delay period) phases of the task was similar or different across dmPFC and OFC. Specifically, we aimed to test the hypothesis that, when activity is averaged across trials, dmPFC and OFC ensembles encode similar task features to similar extents, consistent with a common code for the task.

To do so we calculated the firing rate vector across single units in a brain region during single passes through twenty evenly spaced bins tiling each path (Extended Data Fig. 5a), and similarly through 250-ms time bins tiling the delay period at each well (Extended Data Fig. 5b). We averaged the firing rate vectors in each bin along the same path or at the same well separately for even and odd trials (Extended Data Fig. 5c), yielding one average firing rate vector for each spatial bin along each path on even or odd trials, and likewise for time bins in the delay period at each well. Encoding of the progression along individual paths or through the delay period at individual wells would manifest as greater proximity (see Methods) of average firing rate vectors on even and odd trials at the same bin as compared to across bins. Similarly, if this encoding is shared across paths (for path traversals) or wells (for the delay period), this would manifest as greater proximity within the same bin across sets of trials taken from different paths (for path traversals) or wells (for the delay period), as compared to across different bins (Extended Data Fig. 5d). On the other hand, a proximity value of zero indicates average firing rate vectors are equally close regardless of the rat’s relative distance along paths (for path traversals) or time in the delay (for the delay period).

These analyses of the proximity of trial average population vectors revealed codes for progression towards goals with similar patterns of generalization in the dmPFC and OFC (Fig. 4a–d, Extended Data Fig. 6a,b). During path traversals, dmPFC and OFC average firing rate vector proximity was often above zero both within and across paths (Fig. 4a, Extended Data Fig. 6a), indicating a generalized representation of progression along paths in both dmPFC and OFC. Proximity values were further consistent with a hierarchical code for sensorimotor context along paths, as values were highest along the same path, intermediate along paths through different spaces but sharing the same turn sequence (“same-turn” paths), and lowest along paths through different spaces and with different turn sequences (“different-turn” paths) (Fig. 4a, Extended Data Fig. 6a). Interestingly, average firing rate vector proximity also varied across the span of same- and different-turn paths, with the lowest proximity seen around the turns of different-turn paths (Fig. 4a,c, Extended Data Fig. 6a). In contrast to dmPFC and OFC, HPc average firing rate proximity was high within the same path but very low across paths, highlighting the relative similarity of the dmPFC and OFC codes (Fig. 4a,c, Extended Data Fig. 6a). Altogether, these results indicate that dmPFC and OFC ensembles encode the general progression along a path as well as sensorimotor specifics that distinguish paths through the relative closeness of firing rate states, indicating that the hierarchical code for task progression and context that we previously observed in dmPFC and OFC single unit activity also manifests in population activity.

**Fig. 4.**
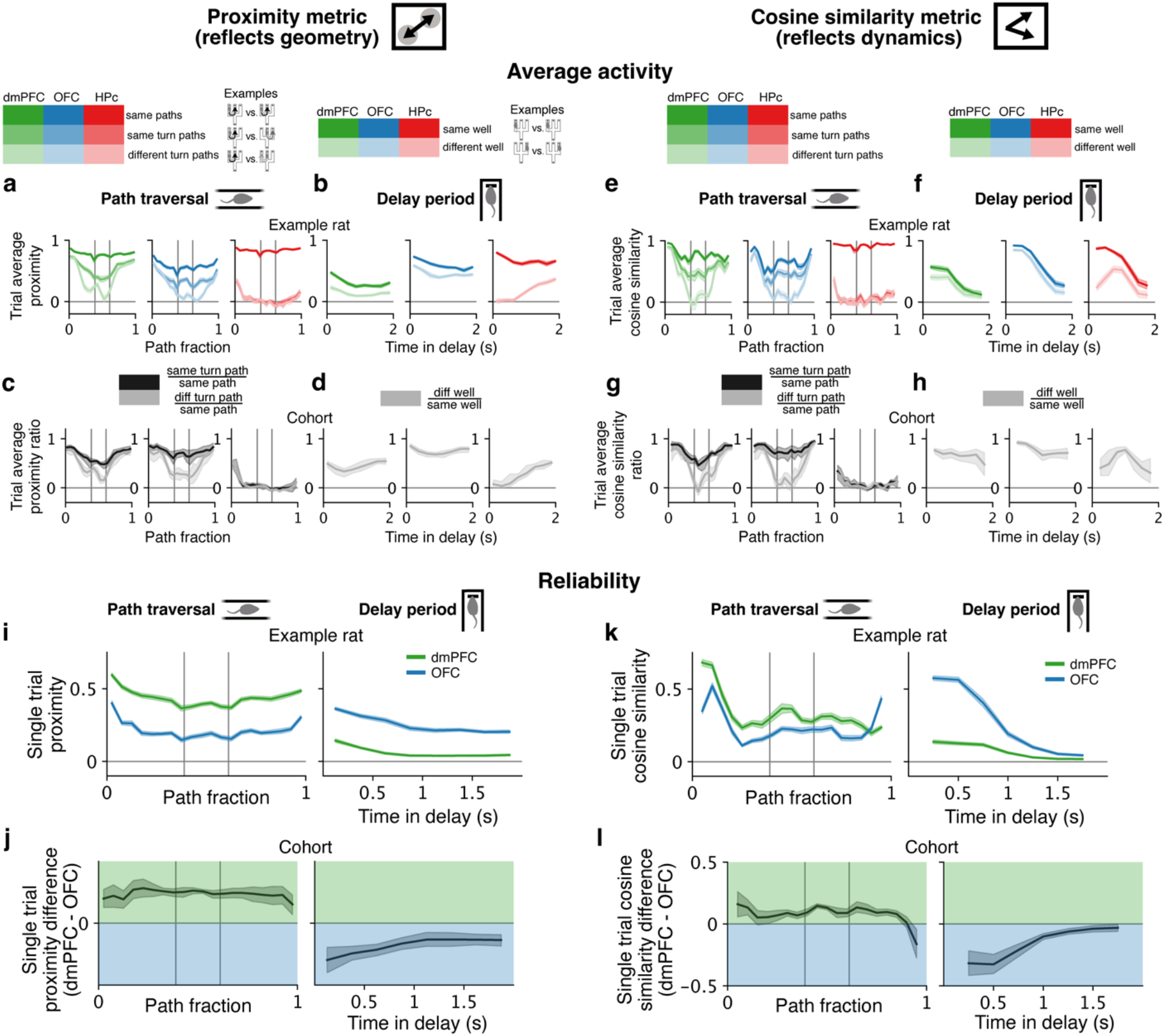
A common code for the task is reliable at different task phases across dmPFC versus OFC ensembles. **a**, Proximity of dmPFC (left), OFC (middle), and HPc (right) average firing rate vectors as a function of fractional distance along the same path (darkest shade), different paths with the same turn sequence (medium shade), or different paths with different turn sequences (lightest shade) (“trial average proximity”) shown for an example rat (mean and 95% CI, *n* = 22 sessions). Vertical lines denote maze junctions. **b**, For the same rat in **a**, proximity of average firing rate vectors as a function of progression through the delay period at the same well (darkest shade) or different wells (lightest shade) (mean and 95% CI, *n* = 22 sessions). **c**, Across rats, ratio of trial average proximity on same turn paths to on the same path (darkest shade), and on different turn paths to on the same path (lightest shade), for dmPFC (left; *n* = 64 sessions from *n* = 3 rats), OFC (middle; *n* = 105 sessions from *n* = 5 rats), and HPc (right; *n* = 83 sessions from *n* = 4 rats) (mean and 95% CI). **d**, Across rats, ratio of trial average proximity during the delay period at different wells to at the same well for dmPFC (left; *n* = 64 sessions from *n* = 3 rats), OFC (middle; *n* = 105 sessions from *n* = 5 rats), and HPc (right; *n* = 83 sessions from *n* = 4 rats) (mean and 95% CI). **e–h**, as in **a–d** but with the cosine similarity of average firing rate difference vectors. **i**, Proximity of dmPFC (green) and OFC (blue) single trial firing rate vectors as a function of path progression (“single trial proximity”) shown for the same rat in **a** (mean and 95% CI, *n* = 22 sessions). **j**, Difference between dmPFC and OFC single trial proximity across rats (mean and 95% CI, *n* = 64 sessions from *n* = 3 rats). **k,l**, as in **i,j** but with the cosine similarity of single trial firing rate difference vectors.

A similar dichotomy between dmPFC and OFC versus HPc was seen during the delay period. During this period, dmPFC and OFC average firing rate vector proximity was above zero both within and across wells (Fig. 4b, Extended Data Fig. 6b), indicating a generalized representation of progression through the delay period in both regions. Proximity values were also consistent with a representation of individual reward well identity, as values were higher at the same versus different wells (Fig. 4b,d, Extended Data Fig. 6b). In comparison, HPc average firing rate vector proximity was relatively higher at the same versus different wells (Fig. 4b,d, Extended Data Fig. 6b).

The analysis of firing rate vector proximity captures one aspect of population activity: the relative distances between firing rate vectors. Intuitively, these distances would be expected to relate to differences in the firing rates of neurons in downstream areas if those downstream neurons are sensitive to the firing rates of their inputs. Population activity can also vary in its dynamics, so we also quantified the direction of evolution of the population activity from one bin to the next and computed the cosine similarity of these firing rate difference vectors, as a measure of the similarity in the direction of progression of population activity (see Methods, Extended Data Fig. 5e). This metric can be thought of as approximating the differences in firing of downstream neurons if those neurons are sensitive to firing rate changes in their inputs.

As with measures of proximity, measures of population dynamics in dmPFC and OFC showed similar patterns of task progression encoding when activity was averaged across trials. Population activity evolved similarly during traversals of different paths and in the delay period at different wells to a greater extent in the dmPFC and OFC than in HPc (Fig. 4e–h, Extended Data Fig. 6c,d). Likewise, the hierarchical code for context observed in dmPFC and OFC firing rate vectors also manifested in the dynamics of those vectors, with higher cosine similarity values along the same path followed by same-turn paths and finally different turn paths during path traversals (Fig. 4e,g, Extended Data Fig. 6c), and likewise at the same well followed by different wells during the delay period (Fig. 4f,h, Extended Data Fig. 6d).

While trial-averaged firing rates provide important insights into overall patterns of activity, they fail to capture the within-trial activity patterns that the brain must itself use during choices and around outcomes. We therefore compared population proximity and dynamics across different trials, asking to what extent neural activity enters similar states and proceeds in similar directions on different passes down the same path or across different delay periods at the same well (see Methods, Extended Data Fig. 5f–h). These analyses quantify the reliability of population-level coding across trials.

The results of these analyses indicate that dmPFC ensembles encode progression towards goals more reliably during path traversals as rats express their choices, whereas OFC ensembles encode progression more reliably as rats arrive at the wells and into the delay period as they wait to learn outcomes. As rats traveled along paths, single trial firing rate vector proximity and dynamics were substantially more consistent across trials in the dmPFC as compared to the OFC (Fig. 4i–l, Extended Data Fig. 7a,b). On the other hand, as rats arrived at reward wells and then waited to learn outcomes, population activity proximity and dynamics were substantially more consistent across trials in the OFC as compared to the dmPFC (Fig. 4i–l, Extended Data Fig. 7a,b). These findings were robust across environments and sets of reward wells (Extended Data Fig. 7d,e). Thus, there is a transition in reliable coding from the dmPFC during path traversal to the OFC as animals arrive at a potentially rewarded place and into the delay before trial outcome is revealed.

We also considered the possibility that the differential reliability we observed reflects reliable coding of unmeasured variables such as motivation, satiation, expectation of future reward, etc., that differed across trials (Extended Data Fig. 5i). To determine whether reliable coding for unmeasured variables could explain our results, we computed the cosine similarity of 100-ms average difference vectors to their nearest neighbor in firing rate space across the entire session (see Methods, Extended Data Fig. 5i) and examined the average of this nearest neighbor cosine similarity in progression bins along paths and in the delay. This assesses whether similar firing rate activity evolves in the same direction across trials, as the encoding of unmeasured task-relevant variables would predict.

Our results were not accounted for by the reliable encoding of unmeasured task-related variables. We found similar patterns of dynamic reliability in dmPFC and OFC as in our prior approach, with dmPFC exhibiting greater nearest neighbor cosine similarity during path traversals, and OFC exhibiting greater nearest neighbor cosine similarity as rats arrived to wells and then waited to learn outcomes (Extended Data Fig. 7c,f). These results suggest that our findings above are best explained as dmPFC and OFC exhibiting reliable patterns of activity at distinct behavioral phases of a cognitive task.

We then validated our findings with a decoding approach, by asking whether the patterns of dynamic reliability we observed in dmPFC and OFC relate to the extent to which task progression can be read out from these areas during distinct task phases. We trained a state space Bayesian decoder on spike time–task progression relationships in 2-ms bins using randomly selected subsets of 50 units in each area, then used the resulting model to predict task progression from spiking in 2-ms bins on held out trials during path traversals or delay periods. Consistent with our single cell and population activity results, task progression could be more accurately decoded from dmPFC activity during path traversals, and from OFC during the delay period (Extended Data Fig. 8a,b).

### Reliable coding relates to future behavior

The distinct patterns of coding reliability across dmPFC and OFC are broadly consistent with their known respective roles in action selection^2,7,8^ and waiting for outcomes^3,9^. We next wondered whether firing rate patterns that differentiate future events related to action selection or waiting for outcomes are expressed by dmPFC and OFC ensembles and if so, whether the reliability of these patterns shows a similar regional specialization. Specifically, we tested the hypothesis that the proximity and dynamics of firing rate vectors that distinguish future behavior are more reliable in the dmPFC during active expression of choices along paths, versus in the OFC as rats wait to learn outcomes. Previous studies have shown activity related to future behavior in both dmPFC and OFC consistent with these regions encoding similar quantities^14,16,33,34^, but whether that activity is more reliable during task periods associated with each region’s critical functions had not been examined.

We began by examining activity as rats traveled along the central stem of the maze from the home well, prior to a left or right choice (“outbound trials”). We compared population activity proximity and dynamics on outbound trials in which rats subsequently made the same versus different choice. Firing rate vector proximity was higher in both dmPFC and OFC on trials with the same future choice, consistent with a signal for upcoming choice (Fig. 5a,b, Extended Data Fig. 9a), but values were relatively higher in the dmPFC (Fig. 5c, Extended Data Fig. 9a). Interestingly, while the same trends were visible for the population dynamics across animals, the differences between dmPFC and OFC were less pronounced, indicating that the similarity of the firing rates across trials was a better predictor of upcoming left or right turn choices than the dynamics of those vectors (Fig. 5f). These findings are consistent with the critical role for the dmPFC in supporting ongoing actions and choices^2,8^.

**Fig. 5.**
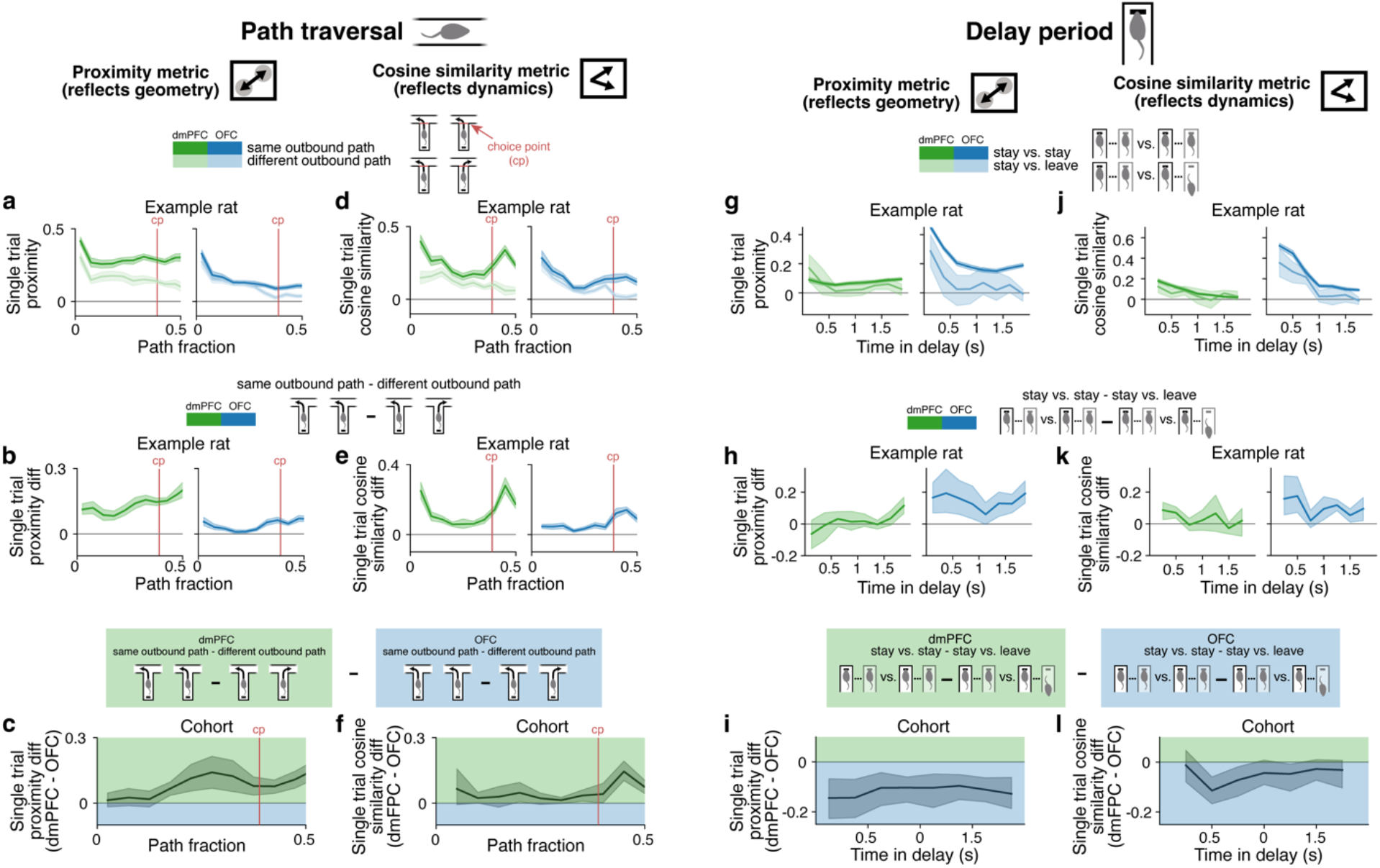
Regional enhancement in the reliable expression of firing rate states that distinguish upcoming events in the dmPFC and OFC. **a**, Proximity of dmPFC (left) and OFC (right) single trial firing rate vectors along the central stem of the maze prior to the choice point (cp) on trials in which the destination well was the same (“same choice”, dark shade) or different (“different choice”, light shade) shown for an example rat (mean and 95% CI, *n* = 21 sessions). In **a–f**, the red line marks the choice point (cp). **b**, For the same rat in **a**, the difference between same choice proximity and different choice proximity in the dmPFC (left) and OFC (right) (mean and 95% CI, *n* = 21 sessions). **c**, Difference between dmPFC and OFC same choice minus different choice proximity across rats (mean and 95% CI, *n* = 64 sessions from *n* = 3 rats). **d–f**, As in **a–c** but with the cosine similarity of single trial firing rate difference vectors. **g**, Proximity of dmPFC (left) or OFC (right) single trial firing rate vectors as a function of progression through the delay period at the same well across “stay” trials (where the rat remained at the well for the full delay period) (dark shade) or across stay and “leave” trials (where the rat left the well prior to the end of the delay period) (light shade) shown for the example rat in **a** (mean and 95% CI, *n* = 21 sessions). **h**, Difference between stay vs. stay proximity and stay vs. leave proximity in dmPFC (left) and OFC (right) shown for the example rat in **a** (mean and 95% CI, *n* = 21 sessions). **i**, The difference between dmPFC and OFC stay vs. stay minus stay vs. leave proximity across rats (mean and 95% CI, *n* = 64 sessions from *n* = 3 rats). **j–l**, as in **g–i** but with the cosine similarity of firing rate difference vectors.

By contrast, lesions to the OFC but not dmPFC have been shown to produce impulsive behavior and impair staying at potentially rewarded sites^3^. In our task, rats occasionally left reward wells before the full delay period had elapsed (173 trials across *n* = 5 rats; average 1.6 trials per session). We therefore compared the reliability of single trial firing rate vector proximity and dynamics in dmPFC and OFC across delay periods where rats stayed at reward wells (“stay-stay”), or across delay periods when animals stayed or left prior to learning the outcome of their choice (“stay-leave”). Only delay times at the well prior to leaving were considered.

We found that reliable coding of progression towards outcomes selectively broke down in the OFC when rats left before outcomes were revealed, consistent with a critical role for the OFC but not the dmPFC in waiting for potential reward^3^. In the OFC but not dmPFC, firing rate vectors were more similar on trials when rats stayed at wells, than on trials when rats stayed versus left (Fig. 5g–i, Extended Data Fig. 9c). And once again, while the same trends were visible in the population dynamics, the differences between OFC and dmPFC were less pronounced (Fig. 5j–l, and Extended Data Fig. 9d) suggesting that the similarity of the firing rates across trials was a better indicator of early leaving behavior than the dynamics of those vectors.

## DISCUSSION

Our findings provide a potential resolution to a long-standing problem in the field: how is regional specialization of function revealed through lesion or inactivation studies consistent with findings that most task variables are encoded across many different brain regions? Here we found that two different regions of prefrontal cortex known to support distinct cognitive functions yet exhibit similar patterns of encoding, dmPFC and OFC, encode similar quantities in rats performing a cognitive task but with dynamic patterns of reliability consistent with specialization for choice- or outcome-related computations.

The common code seen in both dmPFC and OFC represents task progression across multiple levels of generality, capturing the similarities and differences in actions and locations associated with a given trial. These findings are consistent with previous demonstrations of single neuron activity patterns related to progression towards goals^17,22,33,35,36^, turn directions^16,32^, and reward locations^16,33^ in these areas, but precisely how those varying representations are jointly expressed in each region was unclear. Our results identify a similar systematic organization in dmPFC and OFC, with a predominant encoding of the abstract progression towards goals and a simultaneous, less pronounced encoding of the specific spatial and sensorimotor contexts in which those progressions occur.

At the same time, the trial-to-trial reliability of activity and encoding differed profoundly across the two frontal regions, and patterns of reliability matched the previously demonstrated specializations of dmPFC for online decision-making and action^2,3,8^ and OFC for outcome-related computations^2,3,7,9–12^ based on damage to or inactivation of these regions.

Dorsal mPFC exhibited more reliable firing patterns during path traversals. As different dmPFC activity patterns were active at different task phases, this sort of representation is well suited to provide input to motor regions that could directly drive specific action patterns. Consistent with this possibility, dmPFC firing rate states on the central stem were more similar across trials when the future turn choice was the same compared to when it was different. This bias was more pronounced in the dmPFC than OFC, consistent with the critical role for dmPFC but not OFC in action selection^2,7,8^.

By contrast, the OFC exhibited more reliable patterns around the time that animals arrived at reward wells where outcomes are revealed and into the delay period that followed as they waited to learn outcomes. During this period, different OFC activity patterns were active at different times, potentially providing a prediction of when reward would arrive. On trials when rats left reward wells before outcomes were revealed, the reliability of these patterns degraded in the OFC but not dmPFC prior to well departure, consistent with a critical role for the OFC but not dmPFC in waiting for potential reward^3,9^. As OFC activity distinguished different reward wells during the delay period^33^, greater reliability during this period could potentially also prime the OFC or other brain regions for location-specific updates to internal models once outcomes are revealed^2,7,10–12,37^.

We may then ask, why would OFC display unreliable task-related representations during the path traversal and why would dmPFC display unreliable task-related representations during the well arrival and delay? We speculate that these representations reflect the strong interconnectivity of brain regions, where even when a region is not engaged in an ongoing computation, inputs from engaged areas could drive activity in that region. This might also be related to recent studies showing that information about movement is strongly encoded throughout the rodent brain^5,38^. These studies did not assess the relationship between the encoding of movements that are presumably not used to perform a task and the reliable encoding of task-relevant information, and one intriguing possibility is that these are inversely related. Specifically, we might expect that the engagement of task-relevant computations would suppress the expression of task-irrelevant movement representations, whereas movement representations would “leak” into various areas when those areas are not engaged in critical ongoing computations manifesting in more variable activity patterns across single trials.

Our ability to evaluate reliability and compare it across regions was greatly facilitated by simultaneous sampling of large populations of neurons in each region under identical behavioral conditions. These populations enabled us to compute two forms of reliability related to the similarity of population firing rate vectors and the similarity of the changes in those vectors across trials. The large populations also made it possible to determine that our results were unlikely to be the result of reliable encoding of task-related variables that we did not measure.

Our results represent a departure from past ways of thinking about regional specialization from a neurophysiological standpoint, which has historically been to consider the extent to which different variables are encoded in different areas^13–15^. Past studies have identified similar kinds of variables represented across prefrontal regions, either in similar proportions or with regional biases^13,14,24^. Our findings confirm a similar organization of activity but also identify a different axis for evaluating the likely contributions of a brain region to ongoing computations: the trial-by-trial reliability of the activity. This axis identifies choice- and outcome-related periods for dmPFC and OFC as more reliable, respectively, consistent with the lesion and inactivation findings from previous studies^2,3,7,9–12^. We therefore suggest that reliability is a potential signature of when specific brain regions are critical for ongoing computations.

## METHODS

### Animals

Data from five male Long-Evans rats (550–700g) (Charles River Laboratories) were included in this study. Prior to continuous recordings, rats were housed in a temperature- and humidity-controlled facility on a 12-hour light/dark cycle (6AM–6PM). Rats were initially housed with 1– 2 cage mates and had unlimited access to chow. Rats were singly housed once food restriction began. During continuous recordings, rats were housed in a separate facility on a 12-hour light/dark cycle (7AM–7PM). All procedures were approved by the University of California San Francisco Institutional Animal Care and Use Committee.

### Pre-surgical linear track training

Rats were food restricted to 85–90% of their free-feeding weight and trained to run on an elevated linear track (1.1m, 84cm elevation from floor) with reward wells affixed on either end^39^. This training served to familiarize subjects with running on an elevated surface and receiving reward at set locations in an environment. The training also taught subjects to wait up to two seconds at locations to receive reward. In each training session, rats were placed on the linear track for 15 minutes and allowed to behave freely. They were rewarded with a drop of sweetened evaporated milk reward (Nestle sweetened condensed milk plus 25g of sugar; approximately 80– 82μL) at reward wells. Milk reward was delivered automatically via syringe pump through plastic tubing (Tygon, 3.18mm inner diameter, 6.35 outer diameter). Reward became available at a well once rats visited the other well. Over successive training sessions, a temporal delay between when rats arrived to a well and when reward was delivered was gradually introduced. The duration of the delay was fixed within each training session, and increased across training sessions from 0s, to 0.5s, to 1s, to 2s in duration. Once rats achieved a criterion of receiving 30 or more rewards in a session, the delay increased in the next session. Upon reaching this criterion with the 2s delay, pre-surgical linear track training ended, and subjects returned to a diet of ad libitum chow.

### Tetrode microdrive

Tetrodes were spun from 12.5μm diameter nichrome wire and annealed via heat gun using an automated tetrode spinner (SpikeGadgets). Twenty-four tetrodes were loaded into independently moveable shuttles within a 3D printed body^39^.

### Polymer probes

One hundred twenty-eight channel polymer probes were obtained from Lawrence Livermore National Laboratory. Each probe has four shanks spaced 250μm apart. Each shank has 32 linearly arranged contacts (15μm diameter) spaced 26μm apart. Probes were sterilized with ethylene oxide prior to implantation.

### Electrode implantation

Rats underwent stereotactic implantation of 128-channel polymer probes (Lawrence Livermore National Laboratory) and a tetrode microdrive under anesthesia and sterile conditions. Rats were deeply anesthetized with isoflurane, and a mixture of ketamine (50 mg/kg), xylazine (6 mg/kg), and atropine (0.14 mg/kg) was injected intraperitoneally. For the duration of the surgery, anesthesia was maintained using isoflurane and additional intraperitoneal injections of a mixture of ketamine (25 mg/kg) and atropine (0.07 mg/kg) as needed. Body temperature was maintained using a water-based heating pad. Hydration was maintained via subcutaneous delivery of lactated ringers. Lidocaine (0.2 mL) was injected locally into the scalp and an anterior-posterior incision was made to expose the skull. Connective tissue was carefully removed from the surface of the skull. A ground screw was placed over the cerebellum, and additional screws were placed in the rear and front of the skull to provide additional anchoring for the implant.

Polymer probes were targeted to one or both hemispheres of the dorsal medial prefrontal cortex (dmPFC) (+3.2 mm AP, ±0.89–0.94 mm ML, -3.2– 3.6 mm DV, 0–10° tilt; all DV coordinates measured relative to brain surface) and/or orbitofrontal cortex (OFC) (+3.94–4.5 mm AP, ±1.91–2.34 mm ML, -3.6–4.05 mm DV) using a previously published approach^40^. In one animal, polymer probes were targeted to ventral mPFC (+3.2 mm AP, ±0.41–0.44 mm ML, - 4.28–4.33 mm DV) rather than dmPFC. A silicon elastomer (Kwik-Sil) was used to seal craniotomies. A 24-tetrode microdrive was placed over the dorsal hippocampus (-3.8mm AP, ±2.6mm ML). Hippocampal craniotomies were sealed with a silicon sheet. The microdrive was then anchored to the skull using dental cement. A custom-build hybrid headstage (SpikeGadgets) was attached to the electronic interface board on the tetrode microdrive and to the intan boards connected to probes. A 3D printed funnel was placed around the front of the implant and filled with silicon elastomer in order to stabilize the probe electronics. A 3D printed case was placed around the entire implant. Bupivacaine (0.2 mL) was injected locally in the scalp, and sutures were placed to approximate skin in front of and behind the implant. Buprenorphine (0.01–0.02 mg/kg) and meloxicam (2 mg/kg) were administered subcutaneously following surgery for analgesia, and enrofloxacin (5 mg/kg) was given as an antibiotic. Rats were closely monitored in the days following the surgery for signs of discomfort and additional doses of buprenorphine and meloxicam were administered as needed to achieve analgesia.

### Tetrode adjustment

In the weeks following surgery, 1–2 tetrodes were targeted to corpus callosum to serve as a reference, and the remaining tetrodes were slowly advanced to dorsal hippocampus. The presence of sharp-wave ripples and the orientation of sharp waves were used to estimate tetrode depth relative to the hippocampal cell layer^41^.

### Post-surgical linear track training

Between 3 and 6 weeks after surgery, rats were food restricted to 85–90% of their free-feeding weight and reintroduced to running on the linear track. Every 1–2 days for 1–1.5 weeks, rats performed 1–3 15-minute sessions per day on the linear track with a 2s delay between well arrival and reward delivery. During this period, rats were also habituated to the home cage that would serve as their overnight housing during 24-hour recordings.

### Fork maze

Behavior took place in fork-shaped mazes. Each maze was custom-built from acrylic (TAP Plastics). Each maze consists of four arms measuring 73.0 cm attached a central segment measuring 82.2 cm. Two of these arms, the “handle” and “center” arms, attach perpendicularly to the connecting segment at its midpoint and head in opposite directions. The other two arms, the “left” and “right” arms, attach to the connecting segment at its left and right endpoints and head parallel to the center arm. The passageways along the arms and connecting segment are 7.6cm wide and flanked by walls 3.7cm tall. At the ends of each arm is an expanded rectangular segment measuring 12.1cm by 11.1cm and containing a reward well^39^. Each reward well has an opening for milk reward delivery and an infrared emitter and diode to detect arrival to wells. Mazes were elevated approximately 81cm from the floor.

### Alternation task

In the fork maze, rats learned and subsequently performed a memory-based spatial alteration task^42^. In the task, rats receive milk reward at wells according to an alteration rule (Fig. 1b). The rule specifies that subjects should make alternating visits to the left and right wells from a designated home well. Thus, from the left and right wells, rats receive reward at the designated home well, and from the designated home well, rats receive reward at the least recently visited of the left and right wells. On the first well visit of each session, rats receive reward at any of the left well, right well, and designated home well.

An automated behavioral program written in Statescript (Spike Gadgets) detects infrared beam breaks at reward wells and delivers reward via syringe pump through plastic tubing (Tygon, 3.18 mm inner diameter, 6.35 mm outer diameter) following a 2s delay. The syringe pumps were estimated to deliver 81–85 uL of reward, with the exception of one animal for whom pumps were estimated to deliver 64–82 μL of reward due to a pump programming error.

Well visits were deemed “correct”, “incorrect”, or “neither correct nor incorrect” according to the following convention: on trials in which rats could possibly receive reward according to the alternation rule, a well visit is considered correct if rats receive reward and incorrect if they do not. On the other hand, if rats did not have the opportunity to receive reward per the alternation rule, as on the first well visit of a session or when starting from the arm that is never rewarded in the sequence, a well visit is considered neither correct nor incorrect.

### Task environments

Rats initially learned to perform the alternation task in two distinct environments, “Haight left” (HL) and “Haight right” (HR), located within a first room (Room 1) (Extended Data Fig. 1a). Each environment consisted of a dedicated fork maze surrounded by a distinct set of global cues, and walls between environments prevented rats from seeing one environment while located in the other. Rats later learned to perform the task in a third environment, “San Anselmo” (SA), located in a second room (Room 2) (Extended Data Fig. 1a).

### Continuous neural recordings

Throughout behavior on the alternation task, including the initial and all subsequent exposures to the task and the environments, continuous neural recordings were performed 7 days a week and 24 hours a day with the exception of time between recording sessions or in sporadic cases of recording hardware failure. Rats were weighed daily prior to the first sleep session and following the final sleep session and received chow after daily experiments to maintain 85–90% of their free-feeding weight. Occasionally, tetrodes were adjusted after the final sleep session. Rats were housed overnight in a home cage (33 cm x 65 cm x 51 cm) located in Room 1.

### Daily training sessions on the alternation task

Each day, rats performed up to eight 20-min behavioral sessions. In each session, the rat was placed on one of the three fork mazes and allowed to behave freely. Rats were rewarded according to the alternation task as described below. These behavioral sessions were flanked by rest sessions approximately 45-min in length in a small box (32 cm x 33 cm x 41 cm) located in Room 1.

### Learning and performing the alternation task across contexts

Rats underwent a series of exposures to the alternation task in distinct contexts (Extended Data Fig. 1a). These contexts varied in two ways: the environment in which the task was performed, and the identity of the designated home well. Rats initially learned the alternation task in a first environment (E1), either HL or HR, and with a first sequence of well visits (S1), in which the designated home well was either the handle well or center well (Extended Data Fig. 1ai). These assignments were counterbalanced across rats. Once rats performed at least 80% of the total number of correct and incorrect well visits in a session correctly (“≥ 80% correct”) in each of two consecutive sessions, they performed an additional 2–3 sessions at ≥ 80% correct. Rats were then exposed to the alternation task in novel environments and with a novel set of reward wells.

In each novel context subsequently encountered, rats were considered to have reacquired the task once they performed ≥ 80% correct in each of two consecutive sessions. Each exposure to a novel context occurred on a new day, with the exception of a single context exposure for one of the rats. On the day of each novel context exposure, rats first performed “reminder sessions” in one or two familiar contexts. Rats typically performed these sessions at ≥ 80% correct, but in the event they did not, they continued performing sessions in the familiar context until performing ≥ 80% correct.

Rats first reacquired the task in a novel environment (E2) with the previously learned sequence (S1) (Extended Data Fig. 1aii). For each rat, E2 was the environment among HL and HR that had not yet been encountered. Upon performing ≥ 80% correct in the novel context, rats performed S1 in E1 and S1 in E2 on alternating sessions until performing ≥ 80% correct in two consecutive sessions, one in each environment (Extended Data Fig. 1aiii). Rats next learned S2 in E1 (Extended Data Fig. 1aiv). Upon reaching criterion, rats performed S1 in E1 and S2 in E1 on alternating sessions until performing ≥ 80% correct in two consecutive sessions, one on each sequence (Extended Data Fig. 1v). Rats were then exposed to S2 in E2 and performed sessions until performing ≥ 80% correct (Extended Data Fig. 1avi).

Rats then entered a post-acquisition phase, during which they switched among all four sets of environment and sequence according to several rotations (Extended Data Fig. 1avii). Rotation 1 is defined as E2S1, E1S1, E1S2, E2S2. Rotation 2 is defined as E2S1, E2S2, E1S2, E1S1. Rotation 3 is defined as E2S1, E1S2, E2S2, E1S1. Rotation 4 is defined as E2S1, E1S1, E2S2, E1S2. Rats cycled twice through rotation 1, twice through rotation 2, once through rotation 3, and finally once through rotation 4. In this report, we analyzed data from this post-acquisition phase.

Rats subsequently went on to perform the task in additional novel contexts, including a variant in which the designated home well reversed within a single session, as well as in a third environment in a different room (Extended Data Fig. 1avii–xiv). Those data will be analyzed in future reports.

### Histology

Following the final day of recording, rats were deeply anesthetized with isoflurane, and electrolytic lesions were made at the ends of tetrodes. A day or more later, rats were deeply anesthetized with isoflurane, euthanized via injection of 1mL euthasol, and perfused with 4% paraformaldehyde (PFA). The head was placed in 4% PFA. After approximately one day in PFA, the brain was extracted and placed in 4% PFA for an additional day. The brain was then transferred to a sucrose solution for approximately five days for cryoprotection, then sliced in 60–80-micron coronal sections on a cryostat. Immunohistochemistry was performed to facilitate visualization of electrode tracts. Hippocampal sections were stained with 4’,6-diamidino-2-phenylindole (DAPI) and cresyl violet or fluorescent Nissl (NeuroTrace 435/455 Blue, ThermoFisher). OFC and mPFC sections were stained to visualize glial fibrillary acidic protein (GFAP) (primary antibody: mouse anti-GFAP antibody, Sigma-Aldrich; secondary antibody: donkey anti-mouse Alexa 594 antibody, ThermoFisher), and with fluorescent Nissl and DAPI. Stained sections were visualized using fluorescence or light microscopy (Nikon Ti-E Microscope) (Extended Data Fig. 1c). We estimated the locations of electrodes by aligning sections containing tracts to the Paxinos and Watson rat brain atlas (6^th^ edition)^43^.

### Data acquisition

Neural data (electric potential at polymer probe and tetrode contacts relative to a ground screw placed over the cerebellum) and behavioral events (infrared beam breaks at reward wells and syringe pump triggers) were continuously sampled at 30 KHz using a Spike Gadgets acquisition system running Trodes v1.8.2. Video of behavior was obtained from an overhead camera at a rate between 41–50 frames per second. A set of red and green LEDs mounted on the rat’s head was used to track position.

### Position tracking

The centroid of red and green LEDs was estimated using an offline tracking algorithm (SpikeGadgets). Head position measurements were obtained through smoothing and interpolation of these centroids over time.

### Data analysis

All analyses were carried out in the context of Spyglass, the Frank laboratory’s open-source, reproducible data analysis framework (https://github.com/LorenFrankLab/spyglass). This framework includes additional packages such as DataJoint for pipeline management^44^ and SpikeInterface^45^ for spike sorting. Code was written in Python version 3.7.

### Spike sorting

Spike sorting was performed to identify putative spikes of single neurons (“single units”). In preparation for spike sorting, hippocampal tetrode recordings were referenced to a tetrode targeted to corpus callosum and bandpass filtered between 600 and 6000 KHz. Cortical recordings from 128 channel polymer probes were referenced to the median of recording traces on the same shank and bandpass filtered between 300 and 6000 KHz. Putative artifact times were identified as the two milliseconds around times when voltage exceeded 8 standard deviations or 500 mV on over one quarter of channels, and these times were excluded.

We performed spike sorting using MountainSort4, an automated spike sorting algorithm^46^. For hippocampal recordings, units were identified within tetrodes. For 128 channel polymer probe recordings, units were identified in neighborhoods of channels within 115 microns of each other. We performed spikesorting across an interval of time during which the recording appeared stable based upon the amplitudes of identified units. For hippocampal recordings, we performed spikesorting on concatenated run sessions within a day when hippocampal recordings appeared stable on this timescale; this was the case for three subjects. For a fourth subject, we performed spikesorting within each epoch. For 128 channel polymer probe recordings, we performed spikesorting on concatenated run and sleep sessions within a day^46^.

A first round of curation was performed on the units identified by MountainSort4 in an automated fashion using metrics calculated in SpikeInterface^45^. Units with metrics exceeding any of the following thresholds were rejected: a nearest neighbor noise overlap of 0.03, an interspike interval violation ratio of 0.0025, and a time shift in the waveform peak of two samples^45,46^.

A second round of curation was performed on the remaining units to further improve the identification of single units. This round of curation involved visual inspection and had two objectives. First, different units that appeared to correspond to a single neuron whose spikes had been incorrectly split during spikesorting were merged. Second, individual units that appeared to correspond to more than one neuron were excluded. In the case of merging units that appeared to have been incorrectly split, we identified putative instances of two scenarios: the spikes of a single bursting neuron split into two clusters, and the spikes of a non-bursting neuron split into two clusters.

To identify unit pairs putatively corresponding to a single bursting neuron, we visually inspected unit pairs that met three criteria: 1) cosine similarity of average waveforms > 0.6; 2) interspike interval violation ratio less than 0.1; 3) cross correlogram constructed from spikes occurring within 100ms of each other has at least 100 spikes and meets an asymmetry criteria: the ratio of the sum of the density on the side of the correlogram with the greatest density and half the density at zero, to the total density, is greater than 0.6, and 4) the average waveform of the unit that tends to spike first versus second within a 200ms window has the larger peak amplitude. Of these candidate unit pairs, those whose spikes appeared to belong to a single neuron exhibiting successive bursts of spikes with decrementing amplitudes within tens of milliseconds were merged.

Second, to identify remaining cases of unit pairs that appeared to correspond to a single neuron, we visually inspected unit pairs with: 1) similarly shaped waveforms as indicated by a cosine similarity of average waveforms of the two units greater than 0.7; 2) interspike interval violation less than 0.1; 3) cross correlogram constructed from spikes occurring within 100ms of each other has at least 100 spikes; 4) waveform amplitude densities have an overlap^47^ greater than 0.7. Unit pairs that appeared to correspond to a single neuron were merged.

A very small number of units that did not pass these criteria, but appeared to correspond to a single neuron, were also merged. A third and final round of curation was performed on the resulting units using the same procedure employed during the first round of curation. Finally, units for which the maximal amplitude across waveforms occurred on a waveform measured on a channel outside of a target region (dorsomedial PFC including prelimbic and anterior cingulate cortex (ACC), OFC, and dorsal hippocampus) were excluded. Finally, within a session, units with an average firing rate less than 0.1 Hz were excluded. The remaining units were used in downstream analyses. The mean number of units in a recording session for each rat is shown in Extended Data Table 1.

### Data inclusion

In this report, we focus on patterns of activity in dmPFC, OFC, and HPc once rats had learned the task well. We analyzed all sessions in the post-acquisition phase beginning from the first full day of this phase through the final day, with the exception of a session that had large artifacts and a session during which the recording hardware malfunctioned. This amounted to 22, 21, 21, 19, and 22 sessions for rats 1–5. During the post-acquisition phase, rats performed the task well above chance (Extended Data Fig. 1b).

In one rat, the tetrode drive was placed too anteriorly, resulting in a large fraction of tetrodes mistargeted to areas anterior to the HPc. A small number of tetrodes appeared to be in anterior HPc (likely area CA2) but due to concerns about the extent to which this data was representative of HPc, we excluded these putative HPc data. In another rat, we targeted the ventral portion of mPFC. We do not analyze this data here due to the small sample size.

### Statistics

We employed a hierarchical bootstrap procedure^48^ to generate confidence intervals and test for statistical significance. The procedure enables generation of confidence intervals for datasets that consist of repeated measurements at multiple levels, for instance data from multiple sessions from multiple subjects. Importantly, the procedure avoids the high false positive rates that occur if data is instead pooled across levels of a hierarchy and then resampled (e.g. resampling pooled sessions across subjects)^48^. The procedure consists of resampling with replacement at each successive level (e.g. first subjects, then sessions), then recomputing the test statistic of interest. This procedure is repeated several times to produce a “bootstrap distribution” of values that mimics what one may have observed from repeating an experiment several times. The 1 − *α* confidence interval of the test statistic is formed from the *α*/2 and 1 − *α*/2 percentiles of the bootstrap distribution.

To test for a significant difference between two group means, we performed the above hierarchical bootstrap procedure using the difference in group means as the test statistic and drawing one thousand bootstrap samples. A difference was deemed significant if the 1 − *α* confidence interval of the resulting bootstrap distribution did not contain zero. To provide information about the strength of significance, we report the smallest significance level among *α* = 0.05, 0.01, 0.001, 0.0001 at which results were significant. Confidence intervals shown in plots were constructed using a significance level of *α* = 0.05.

### Definition of well arrival and departure

Arrival to a reward well was identified as the start of the first “down” state of the digital input detecting infrared beam breaks at the reward well, following a down state at another reward well. Departure from a reward well was identified as the end of the final down state at the location, prior to a down state at another reward well.

### Trials

We define a trial as the time from the departure at one well to the departure at another well. We consider subsets of trials that correspond to distinct behavioral periods, as described in *Task periods*. When clear from context, we refer to these subsets as trials.

### Task periods

For the purposes of analysis, we divided each trial into three task periods. The “path traversal” period begins upon a subject’s departure from a reward well and ends upon their arrival at a different reward well. The “delay” period begins upon arrival to a reward well and ends 2s later. At this time, a milk reward is delivered if the well visit was correct. On rare occasions, a delay period was ongoing at the time the session ended (0.1% of all delay periods); these were excluded from analysis. In 2.3% of the remaining delay periods (173 delay periods; mean per session: 1.6), the rat left the well before the 2s had elapsed; we refer to these as “leave”. Conversely, we refer to delay periods where rats remained at the well for the entire delay period as “stay”. Analyses of delay periods include only “stay” trials unless otherwise noted. The “post-delay” period begins at the end of the delay period and ends when a subject departs from the well.

### Task progression

During path traversals, we define task progression as the fraction of the path rats have traversed. During the delay period, we define task progression as the time elapsed following arrival at a reward well. During the post-delay period, we define task progression as the relative time between the end of the delay period and departure from the reward well.

### Spike rasters

In Fig. 1e, simultaneously recorded units from each brain region were sorted in ascending order by the location of average peak firing during a concatenation of the average firing rate in a session during traversals along the shown path, delay periods at the shown well, and post-delay periods at the shown well.

### Firing rates

Firing rates were estimated by convolving spike times of each single unit with a Gaussian kernel with a standard deviation of 100ms and sampling the resulting vectors every 100ms.

### PCA of population activity

Principal component analysis (PCA) was performed on vectors of firing rates formed from all single units during a behavioral session (Extended Data Fig. 2). For each brain region, the smallest number of principal components (PCs) needed to explain at least 80% of the variance in the data was identified. To assess how this estimate of population activity dimensionality varies with ensemble size, we recomputed the number after randomly subsampling units without replacement to obtain ensemble sizes ranging from 5 to the number of units recorded in the brain region (Extended Data Fig. 2b).

### UMAP embeddings

We used Uniform Manifold Approximation and Projection (UMAP) to visualize two-dimensional non-linear embeddings of the high-dimensional neural activity space in which each axis corresponds to the firing rate of a single unit^27^. UMAP embeds high-dimensional data in a lower dimensional space with the goal of preserving the local distance relationships between nearby data points. We embedded firing rate vectors formed from all single units during individual behavioral sessions.

We colored embedded neural states to visualize their organization with respect to task variables of interest, including task progression (Fig. 2a,b, Extended Data Fig. 3) and path identity (Fig. 2c, Extended Data Fig. 3) during task periods. During path traversals, path identity referred to the path the rat was on. During the delay and post-delay periods, path identity referred to the path the rat took to the current well. We made UMAP embeddings for each brain region for which there were at least 50 single units in a session (Extended Data Fig. 3).

### Estimating the reliability of single unit task progression representations at distinct levels of generality

Progression through task periods occurs within distinct contexts provided by the different paths and reward wells in a maze. In principle, single unit representations of these progressions may be similar ("generalize”) across any number of the distinct contexts. For instance, a single unit may exhibit the same pattern of firing rate modulations over the course of the delay period across several reward wells. We use the term “level of generality” to refer to the number of contexts across which representations of task progression are invariant. Given *N* contexts, there are *N* possible levels of generality (1, …, *N*). During the path traversal period, we consider potentially rewarded paths in a session (*N* = 4), and during the delay period, we consider potentially rewarded wells in a session (*N* = 3). At a given level of generality *k*, the total number of ways that activity can be invariant across contexts is given by the binomial coefficient:

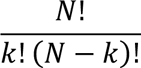

We aimed to estimate the reliability of single unit firing rate patterns that encode task progression at different levels of generality. At a high level, our approach relies on the idea that if the firing of a single unit reliably represents task progression in the same manner across a subset of contexts, then a model of the activity trained in any context in the subset should perform equally well in predicting activity in the other contexts in the subset, and perform poorly in predicting activity in contexts outside the subset. From the observed pattern of cross-context model performance, we can then estimate the reliability of task progression representations at distinct levels of generality.

### GLM of single unit firing in relation to task progression

We modeled single unit firing as a function of task progression using an established GLM framework^49–52^. In the framework, spikes are assumed to be derived from a Poisson process with a time-varying rate parameter that is a function of a linear combination of covariates at each time. The function is chosen to be the exponential function with base *e*, which serves to guarantee that the rate parameter will be positive, as is required for the Poisson distribution. The rate parameter λ is thus:

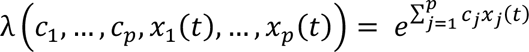

where *x_j_*(*t*) is the j^th^ covariate at time t and *c_j_* is the coefficient for the j^th^ covariate.

We built separate GLMs to predict firing rates during the path traversal and delay periods. We defined sets of covariates to capture task progression in each case using a raised cosine basis^51^. Compared to an indicator basis that represents the presence or absence of the rat in task progression bins using a binary variable, the raised cosine basis allows us to model task progression with relatively fewer covariates, which is expected to reduce overfitting. For GLMs of task progression during the path traversal period, we defined fourteen evenly spaced raised cosine bumps along each of the four potentially rewarded paths, for a total of 56 covariates. For GLMs of task progression during the delay period, we defined fourteen evenly spaced raised cosine bumps across the time in the delay at each of the three potentially rewarded wells, for a total of 42 covariates.

We trained and tested GLMs using the statsmodels python package^53^. During training, Broyden–Fletcher–Goldfarb–Shanno optimization^54^ was used to find coefficients *c*_1_, …, *c*_*p*_ that minimize the following cost function given a set of spike counts and covariates in 20-ms time bins:

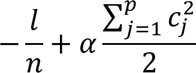

In the first term, *l* is the model log likelihood and *n* is the number of observations, which in our case is the number of time bins in the training set. The second term is an L2 penalty included to improve stability during training. In this term, *c_j_* is the coefficient for the j^th^ covariate, and *α* is a free parameter that scales the impact of the regularization term on the overall cost. We set *α* = 0.00005. During testing, a trained model was used to predict spike counts given covariate values in 20-ms time bins.

### Training and testing GLMs across contexts

We trained a model in each context (paths for path traversals and wells for delay periods) and tested that model in the same and every other context (Extended Data Fig. 4a). For computational tractability, we considered only the subset of contexts that could yield reward according to the variant of the alternation task operative in a given session (“potentially rewarded”).

These analyses were performed across data from many trials, where each trial corresponds to a single traversal of a path in the case of the path traversal period, and a single pass through the delay period at a well in the case of the delay period. For training and testing in the same context, we predicted firing in each trial using a model trained on all other trials (“leave one out cross validation”), then combined results across trials. For training and testing in different contexts, we used all trials in the different context for training.

For each pair of train and test context, we quantified model performance as the Pearson correlation coefficient between predicted and actual firing rates (Extended Data Fig. 4b). We chose this metric as it in part reflects the informativeness of single unit firing about progression through the task period. As an extreme example of this, a cell that fires during a task period but exhibits no reliable modulations will have a low correlation value. By contrast, an area-based measure would yield a high value for such a cell. With regard to our dataset, this is particularly relevant for prefrontal cortex neurons as these tend to have nonzero baseline firing rates, resulting in high area-based values even when no significant modulations in firing rate are present. To obtain firing rates, we convolved spike counts with a 100-ms Gaussian kernel.

### Estimating reliability at distinct levels of generality

For each single unit, we approximated the vector of model performance values for pairs of train and test contexts as a positive linear combination of basis vectors that together capture all possible subsets of contexts (specific paths in the case of path traversals, and specific wells in the case of the delay period) across which single unit firing rate patterns could be invariant (Extended Data Fig. 4c):

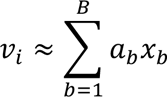

where *v_i_* is the observed vector of correlation coefficients between actual and predicted firing rates for each possible pair of train and test contexts for unit *i*, *x*_*b*_ is the vector of correlation coefficients between actual and predicted firing rates for each possible pair of train and test contexts given generalization across the b^th^ subset of contexts, *a_b_* are positive coefficients, and *B* is the total number of basis vectors. The total number of basis vectors is the sum of the number of ways for firing rate patterns to be invariant across contexts at each level of generality *k*:

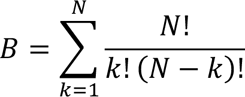

where *N* is the total number of contexts.

Each basis vector contains model performance values that would be expected given a perfectly reliable representation of task progression (i.e. identical pattern of firing rate modulation across trials) that generalizes across a particular subset of contexts. Specifically, a basis vector that reflects generalization of a perfectly reliable task progression representation across a particular set of contexts would have model performance values of one for each train/test pair formed from contexts in the set, and model performance values of zero for all other train/test pairs.

We approximated *v_i_*, the observed vector of correlation coefficients between actual and predicted firing rates for pairs of train and test contexts for single unit *i*, as a positive linear combination of basis vectors using non-negative least squares. This produces a set of weights that reflect the extent to which each basis vector contributes to the observed vector of correlation coefficients. To enable a fair comparison of the weights for basis vectors at different levels of generality, we scaled each weight by multiplying by the number of contexts (paths or wells) to which its corresponding basis vector applies and dividing by the total number of contexts with at least one spike. The upshot of this is that single units that fire perfectly reliably in all contexts in which they are active have a weight sum equal to one. We refer to these scaled weights as “reliability metrics”.

To illustrate this scaling, consider a single unit that fires the same way across trials within each path, but that has uncorrelated firing rate patterns across different paths. Now consider a second single unit that fires the same way across trials across all four paths. The first single unit would have a weight equal to one for each of the basis vectors corresponding to the unique representation of a single path, reflecting that it fires reliably along each path, and the sum of its weights would be four. On the other hand, the second single unit would have a weight of one for the single basis vector that corresponds to generalized representation across the four paths, and the sum of its weights would be one. After scaling these weights by the number of paths to which they apply (one for the first single unit, and four for the second single unit), the two single units would both have a weight sum of one, reflecting that both reliably represent task progression along all paths.

Prior to estimating weights, we set the prediction value to zero for pairs of train and test contexts for which no spikes occurred in the train context (path or well), to reflect the fact that no information about the testing context is available from the training context. Similarly, for pairs of train and test contexts for which no spikes occurred in the test context, we set the prediction value to zero to reflect the fact that there are no events in the test context to predict.

We next aimed to compare the reliability of task progression representations at different levels of generality across brain regions. We first asked whether single units in a brain region tended to express reliable firing rate patterns unique to one path or generalized across, two, three, or all four paths during the path traversal period, or likewise unique to one well or generalized across two or three wells during the delay period. Separately for the path traversal and delay periods, for each single unit in a brain region we summed the reliability metrics at each level of generality. Then at each level of generality we found the fraction of single units for which the summed reliability metrics at that level was greatest (mean and 95% confidence interval across rats and sessions) (Fig. 3g,h). We next asked how the magnitude of reliability compared across brain regions for representations at different levels of generality. Separately for the path traversal and delay periods, for each level of generality we found the mean reliability metric sum at that level across single units for which the sum at that level was greater than at any other level (mean and 95% confidence intervals across rats, sessions, and single units) (Fig. 3i,j).

### Single-trial firing rate maps

In Fig. 3a–c, we computed single-trial firing rates as a function of path fraction in 20 bins spanning the path (approximately 3.65 cm bin width) and convolved the resulting values with a Gaussian kernel of the same width. In Fig. 3d–f, we computed single-trial firing rates as a function of time in the delay period in 100ms bins and convolved the resulting values with a 100-ms Gaussian kernel.

### Proximity and dynamics of neural population activity

We studied population neural activity by considering the high-dimensional space in which each axis corresponds to the firing rate of a single unit in a brain region. At each point in time, the population activity exists at a single point in this firing rate space. We considered two aspects of how population activity evolved in this space during task performance. First, we considered how close firing rate vectors tended to be at the same versus different point in task progression (“proximity”), which relates to their geometry. Second, we considered the extent to which firing rate vectors evolved in similar or different directions at a given point in task progression, which relates to their dynamics. Since the dimensionality of the space can affect these measurements, we performed these analyses in a 50-dimensional space by using random samples of 50 single units recorded in a brain region. We chose a 50-dimensional space as most recorded ensembles in a brain region had at least fifty units. As recorded ensembles could have more than 50 units, we used ten random samples of units from each ensemble when estimating confidence intervals.

We examined firing rate vector evolutions on average to assess average characteristics of population activity as well as on single trials to assess reliability. In both cases, we began by constructing an approximation of the evolution of neural activity on single trials using a piece-wise linear curve^55,56^. On single trials, we defined the knots of the curve as the average firing rate vector across simultaneously recorded single units in a brain region during consecutives samples within each of twenty spatial bins spanning a path (path traversals) (Extended Data Fig. 5a), and eight temporal bins spanning time in the delay at the well (delay period) (Extended Data Fig. 5b). The number of spatial and temporal bins was chosen so that on average, the average time rats spent within a bin was similar across path traversals and the delay period, and so that there were sufficient samples in each bin. The series of these knots over time and the line segments that connect them comprised the curve.

### Average neural population activity trajectories

We found the average of these single trial firing rate vector trajectories by computing the mean firing rate vectors within each path or delay progression bin across trials within a session (Extended Data Fig. 5c). We did so on even or odd numbered trials to enable comparison of average activity within the same path or at the same well. We then compared average activity for sets of conditions that differ in the extent of shared contextual features provided by physical space along paths or at wells, and requirement for similar or different motor actions via shared or different sets of turns along paths. For path traversals, we compared even and odd trials on the same path (same physical space and same turn sequence), even and odd trials along different paths with the same turn sequence (different physical space but same turn sequence), and even and odd trials along different paths with different turn sequences and without overlapping track segments (different physical space and turn sequence). For delay periods we compared even and odd trials at the same well (same physical space) or different wells (different physical space).

### Average proximity of neural population activity

We quantified the extent to which average population activity vectors were relatively closer together at similar versus different points in task progression using a proximity metric (Extended Data Fig. 5d). At each task progression bin, we computed the mean distance of the average firing rate vectors on even and odd trials (Extended Data Fig. 5d), to obtain a measure of the distance between average firing rate vectors from the same task progression bin. At each task progression bin, we also computed the mean distance of the average firing rate vector in that bin on even trials to the average firing rate vector in a different task progression bin on odd trials. This captures the average distance between average firing rate vectors from different task progression bins. We then found the ratio of these average distances. We defined the proximity metric as one minus this ratio, such that larger numbers indicate population vectors that are relatively closer at the same versus different relative distance along paths during the path traversal period, or time elapsed since well arrival during the delay period. Thus, the proximity at task phase bin *i* in context *a* in relation to context *b* is defined as:

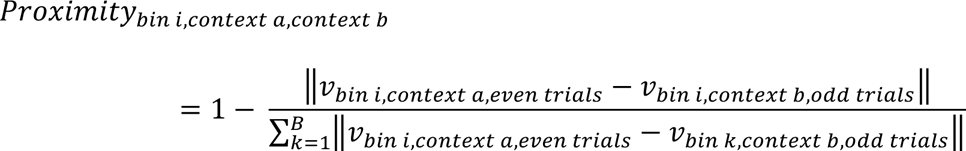

where *v_bin i,context a,even trials_* is the average across even trials of the firing rate vector at task phase bin *i* in context *a*; *B* is the total number of task phase bins; and contexts are paths during the path traversal period and wells during the delay period.

For path traversals, we computed the proximity metric in three cases: using even and odd trials along 1) the same path, 2) different paths with the same turn sequence, and 3) different paths with different turn sequences and without overlapping track segments. For delay periods, we computed this metric in two cases: using even and odd trials at 1) the same well and 2) different wells. We summarized the results for each of these cases for individual rats, by computing the mean and associated 95% confidence intervals of the proximity metric across sessions, pairs of paths (for path traversals) or wells (for delay periods), and ten random samples of 50 single units (Fig. 4a,b and Extended Data Fig. 6a,b).

We then directly compared these cases. For path traversals, we divided the metric obtained using even and odd trials on different paths with the same turn sequence by the metric obtained using even and odd trials on the same path. Likewise, we divided the metric obtained using even and odd trials along different paths with different turn sequences and without overlapping track segments by the metric obtained using even and odd trials on the same path. For delay periods, we divided the metric obtained using even and odd trials at different wells, by the metric obtained using even and odd trials at the same well. We performed these divisions within each session and computed the mean and associated 95% confidence intervals of the resulting ratios across rats, sessions, pairs of paths (for path traversals) or wells (for delay periods), and ten random samples of 50 single units (Fig. 4c,d).

### Average dynamics of neural population activity

We also quantified the extent to which firing rate vectors evolved in a similar direction from one task progression bin to the next on average (dynamics) (Extended Data Fig. 5e). We repeated the procedure above using the cosine similarity of the difference of consecutive average firing rate vectors on even and odd trials (Fig. 4e–h). For each pair of consecutive task phase bins, we defined the cosine similarity of the average difference vectors on even trials in context *a* and odd trials in context *b* as:

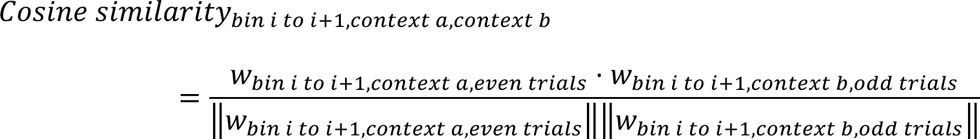

where *w_bin i to i+1,context a,even trials_* is the difference vector spanning bins *i* to *i* + 1 in context *a* on even trials; ⋅ is the dot product; || || denotes vector length; and contexts are paths during the path traversal period and wells during the delay period.

### Reliability of neural population activity proximity

To quantify the reliability of firing rate states across trials (Extended Data Fig. 5f), we computed the Euclidean distance between single trial firing rate vectors separated by at least 10 s that either occurred within the same task progression bin or within different task progression bins along a given path (path traversal) or at a given well (delay period) (Extended Data Fig. 5g). This separation effectively ensured that we compared firing rate vectors from different trials and was designed to minimize the extent to which short-timescale fluctuations in activity that are unrelated to task progression contribute to the measured proximity. We used the same proximity metric defined above to compare the distances of single trial firing rate vectors within the same versus different task progression bin. A greater value indicates that single trial firing rate vectors tend to be closer in the same versus different task progression bins. For each rat and brain region, we found the average of this proximity metric and associated 95% confidence intervals across sessions, pairs of paths (for path traversals) or wells (for delay periods), and ten random samples of 50 single units (Fig. 4i, Extended Data Fig. 7a). We also report the average of this metric within individual sessions, averaged across pairs of paths (for path traversals) or wells (for delay periods) and ten random samples of 50 single units (Extended Data Fig. 7d). To directly compare reliability across dmPFC and OFC, we computed the difference of the proximity metric for dmPFC and OFC in individual sessions for rats with dual dmPFC and OFC recordings. We then found the average of this proximity metric difference and associated 95% confidence intervals across rats, sessions, pairs of paths (for path traversals) or wells (for delay periods), and ten random samples of 50 single units (Fig. 4j).

### Reliability of neural population activity dynamics

To quantify the reliability of firing rate state dynamics across trials, we computed the cosine similarity between single trial firing rate difference vectors in the same task progression bin along each path and at each well in the delay period separated by at least 10 seconds (Extended Data Fig. 5h). We found the average of this measure and associated 95% confidence intervals for each rat and brain region across sessions, pairs of paths (for path traversals) or wells (for delay periods), and ten random samples of 50 single units (Fig. 4k, Extended Data Fig. 7b). We also report the average of this metric for individual rats and sessions, averaged across pairs of paths (for path traversals) or wells (for delay periods) and ten random samples of 50 single units (Extended Data Fig. 7e). We also computed the difference of this metric for dmPFC and OFC in individual sessions for rats with dual dmPFC and OFC recordings. We found the average of this proximity metric difference and associated 95% confidence intervals across rats, sessions, pairs of paths (for path traversals) or wells (for delay periods), and ten random samples of 50 single units (Fig. 4l).

### Reliability of neural population activity dynamics in local regions of neural population activity space

We aimed to understand whether the dynamic patterns of consistent population activity proximity and dynamics we observed are likely to reflect a global disengagement of brain regions during distinct task phases, wherein variables are unlikely to be reliably coded by these regions. Alternatively, periods of unreliable task progression coding could be times when the population is encoding other task features that vary across trials along the same path or in the delay period at the same well (Extended Data Fig. 5i). In this case, we expect neural activity dynamics to be highly similar to those with similar firing rate states (Extended Data Fig. 5i). To assess this, we found firing rate vectors in 100-ms bins spanning an entire session. For the firing rate vector at each time point, we found the firing rate vector at a different time point separated by at least 10 seconds that had the smallest Euclidean distance to the first vector (“nearest neighbor”). We then computed the cosine similarity of the difference vectors at the two time points. We found the average of these “nearest neighbor cosine similarity” values within each path bin and each delay progression bin. For each rat and brain region, we found the average of these values and associated 95% confidence intervals across sessions and ten random samples of 50 single units (Extended Data Fig. 7c). We also report the average values for individual rats and sessions, averaged across pairs of paths (for path traversals) or wells (for delay periods) and ten random samples of 50 single units (Extended Data Fig. 7f).

### Decoding task progression from spiking activity

We decoded task progression during path traversals or the delay period using a previously published Bayesian state space model^57^. In each behavioral session, we defined a set of contiguous 2-ms time bins spanning the session, denoted as *t*_1_, …, *t*_*T*_ where *t*_*k*_ is the k^th^ time bin. We define latent variable *x*_*k*_ that corresponds to the neural representation of task progression at *t*_*k*_.

We define *I_k_*, a discrete latent variable that denotes whether the neural representation of task progression evolves according to continuous or fragmented dynamics. The model simultaneously estimates the posterior probability of task progression and the dynamics: *p*(*x*_*k*_, *I_k_* | *O*_1:*T*_) where *O*_1:*T*_ is the set of observed spike counts in each time bin from 1 through T. The estimation is carried out via a pair of equations. First, we apply a causal filter equation, beginning with initial conditions *p*(*x*_0_, *I*_0_) and recursively iterating from *t*_1_ to *t*_*T*_:

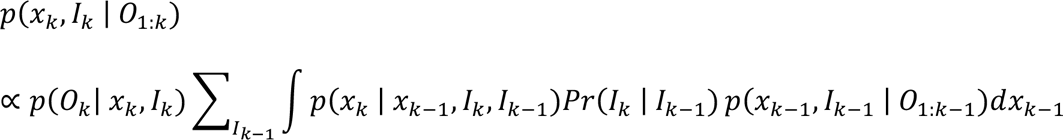

Second, we apply an acausal smoother equation, recursively iterating backwards in time from *t*_*T*_ to *t*_1_:

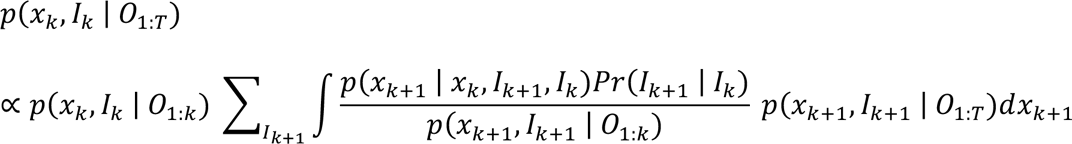

where:

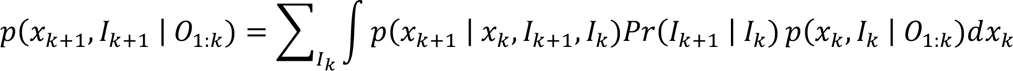

Four quantities must be defined or estimated in order to specify this model: the initial conditions *p*(*x*_0_, *I*_0_), the dynamics movement model *Pr*(*x*_*k*_ | *x*_*k*-1_, *I*_*k*_, *I*_*k*-1_), the dynamics transition matrix *Pr*(*I*_*k*_ | *I*_*k*-1_), and the likelihood of the observations *p*(*O_k_* | *x_k_*, *I_k_*). To reflect our lack of prior knowledge about the initial latent positions and dynamics, we define the initial latent positions and initial dynamics as uniformly distributed:

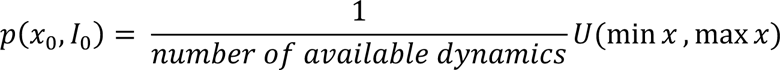

where *U* denotes the uniform distribution.

The neural representations may evolve with distinct dynamics. The algorithm aims to model this explicitly. Two dynamics are specified that respectively capture continuous and non-continuous evolution of task progression. In the “continuous” dynamic, the next latent variable value is normally distributed around the current latent variable value. In the “fragmented” dynamic, the next latent variable value is uniformly distributed over all possible values of the latent variable. We assume that we have no information about the value of the latent variable when transitioning to or from the fragmented state, which we capture using the fragmented dynamic. These choices are reflected in the dynamics movement model:

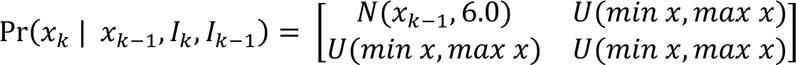

with row indices corresponding to *I*_*k*-1_ = [*continuous, fragmented*] and column indices corresponding to *I_k_* = [*continuous, fragmented*] and where *U*(*left, right*) represents a uniform distribution spanning the left and right limits specified. We define the dynamics transition matrix to reflect the prior expectations that dynamics last 100ms on average, and that there is a small probability of changing to other dynamics:

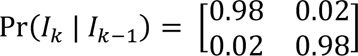

with row indices corresponding to *I*_*k*-1_ = [*continuous, fragmented*] and column indices corresponding to *I_k_* = [*continuous, fragmented*].

We compute the likelihood of observations *p*(*O_k_* | *x*_*k*_, *I_k_*) using an encoding model:

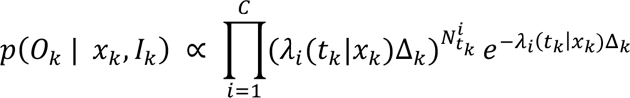

where 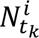 represents a spike in time bin *k* from cell *i*, Δ_*k*_ is the size of the time bin, and *λ_i_*(*t_k_*|*x_k_*) is the instantaneous firing rate of cell *i* given *x_k_*. We estimate *λ_i_*(*t_k_*|*x_k_*) through kernel density estimation.

We applied the above algorithm to decode task progression during path traversals, and separately during the delay period. We decoded path traversals using 100 bins and the delay period using 50 bins. We used a greater number of bins for path traversals because these span a longer period of time than the delay period. In order to use the same parameters across models, we multiplied each variable by a constant factor prior to decoding, then multiplied by the inverse to display results on the original scale.

We performed decoding using leave one out cross validation on trials. Specifically, we trained a model on all but one of the trials on a given path in the case of decoding path traversals, or in the subsequent delay period in the case of decoding time in delay, then tested the model on the held-out trial. We did this for each path in the case of decoding path traversals, and for the delay period following traversals of each path in the case of decoding time in the delay period.

We defined the decoding error as the mean absolute difference between the actual task progression, and the most likely estimate of task progression (maximum a posteriori estimate). To quantify decoding error at different parts of the task, we found the mean decoding error in ten evenly spaced bins spanning path fraction, and in ten evenly spaced bins spanning the delay period, within each recording session. For individual rats, we estimated the mean and associated 95% confidence intervals of these values across sessions, paths (for path traversals) or proceeding paths (for delay periods), and ten random samples of 50 single units (Extended Data Fig. 8a). To determine whether there are differences in decoding error at particular parts of the task when using dmPFC versus OFC single units, we also found the difference of dmPFC and OFC errors in each task progression bin for rats with dual dmPFC and OFC recordings. We found the mean and associated 95% confidence intervals of these values across rats, sessions, paths (for path traversals) or proceeding paths (for delay periods), and ten random samples of 50 single units (Extended Data Fig. 8b).

### Reliability of population activity prior to left vs. right choices during path traversals

We asked whether population activity encoding upcoming choices was more reliable across trials in dmPFC as compared to OFC, consistent with the critical role for the dmPFC but not OFC in action selection^2,7,8^. To test this, we asked whether dmPFC firing rate vectors tended to be closer together and evolve in similar directions as rats approached the choice point from the home well on trials with the same future left or right choice in comparison to on trials with different future choices. In each session, we computed the single trial firing rate vector proximity and single trial firing rate difference vector cosine similarity in each spatial bin along the home arm for pairs of trials with the same destination well (“same outbound path”), and separately for pairs of trials with different destination wells (“different outbound path”). We show the mean and associated 95% confidence intervals of the resulting metrics across sessions, pairs of paths, and ten random samples of 50 single units (Fig. 5a,d, Extended Data Fig. 9a,b).

In each session, we also computed the difference between these metrics across the “same outbound path” and “different outbound path” conditions, as a measure of how much more reliable single trial firing rate vectors and dynamics were on trials when rats made the same versus different choice (Fig. 5b,e). Finally, in each session, we computed the difference of this measure across dmPFC and OFC for rats with dual dmPFC and OFC recordings. This captures the extent to which firing rate vectors and dynamics were more reliable on trials when rats made the same versus different choice in the dmPFC as compared to the OFC. We show the mean and 95% confidence intervals of this difference across rats, sessions, pairs of paths, and ten random samples of 50 single units (Fig. 5c,f).

### Reliability of population activity on stay vs. leave trials during the delay period

Rats with lesions to the OFC are more likely to leave prior to delivery of a reward^3^. We therefore asked whether OFC firing rate vectors tended to be closer together and evolve in similar directions in the delay period at a given reward well on trials when rats stayed to learn the outcome their choice in comparison to on trials when they left before outcomes were revealed, thereby distinguishing upcoming staying and leaving behavior. In each session, we computed single trial firing rate vector proximity and single trial firing rate difference cosine similarity in each temporal bin during the delay period for pairs of trials during which rats stayed at a well to learn outcomes (“stay vs. stay”), and separately on trials where rats stayed versus left a well prior to outcomes being revealed (“stay vs. leave”). Only time bins prior to well departure, when rats were still at the well, were included. We show the mean and associated 95% confidence intervals of the resulting metrics across sessions, pairs of paths, and ten random samples of 50 single units (Fig. 5g,j, Extended Data Fig. 9c,d). In each session, we also computed the difference between these metrics across the “stay vs. stay” and “stay vs. leave” conditions, as a measure of how much more reliable firing rate state and dynamics were on trials when rats stayed, versus across trials when rats stayed versus left prior to learning outcomes (Fig. 5h,k). Finally, in each session, we computed the difference of this measure across dmPFC and OFC for rats with dual dmPFC and OFC recordings. This captures the extent to which firing rate vectors and dynamics were more reliable on trials when rats stayed as compared to trials when rats stayed versus left prior to learning outcomes in the OFC as compared to the dmPFC. We show the mean and 95% confidence intervals of this difference across dmPFC and OFC across rats, sessions, pairs of paths, and ten random samples of 50 single units (Fig. 5i,l).

## Data availability

The datasets analyzed during the current study will be made publicly available in the Distributed Archives for Neurophysiology Data Integration (DANDI) Archive. Any additional information needed to analyze the data reported in this paper is available from the corresponding authors upon request.

## Code availability

All code used for the analysis in this paper will be made publicly available at https://github.com/LorenFrankLab.

## Acknowledgments

We thank Viktor Kharazia and Anya Kiseleva for their assistance with histology; Rhino Nevers and Ryan Ly for their contributions to curation and preprocessing pipelines; Joni Wallis and Karunesh Ganguly for helpful discussions; and Brett Mensh, Shih-Yi Teng, and other members of the Frank laboratory for their input on the manuscript. J.A.G. was supported by National Institute of Mental Health grant F30MH126483, National Institutes of Health grant T32GM007618, UCSF Discovery Fellowship, and Phi Beta Kappa Graduate Award. A.E.C. was supported by NSF GRFP award 1650113, NIH award F31MH124366, and UCSF Discovery Fellowship. A.J. was supported by the Life Sciences Research Foundation grant. K.H.L. was supported by the Helen Hay Whitney Foundation. L.M.F was supported by a Howards Hughes Medical Institute Investigator Award and the Simons Collaboration on the Global Brain grant 521921.

## Contributions

J.A.G. and L.M.F. conceptualized and designed the study. J.A.G. performed surgeries with assistance from D.P.G., A.E.C., and A.J. J.A.G. and D.P.G. collected data. J.A.G. analyzed the data. A.E.C., J.Z., P.T., J.S., A.Y., and R.H. contributed to the development of surgical techniques and equipment. E.L.D. contributed to the development of analytical techniques for decoding analyses. K.H.L. contributed to data preprocessing pipelines. C.K. contributed useful discussion. J.A.G. and L.M.F. wrote the manuscript with input from all authors.

---

**Extended Data Fig. 1.**
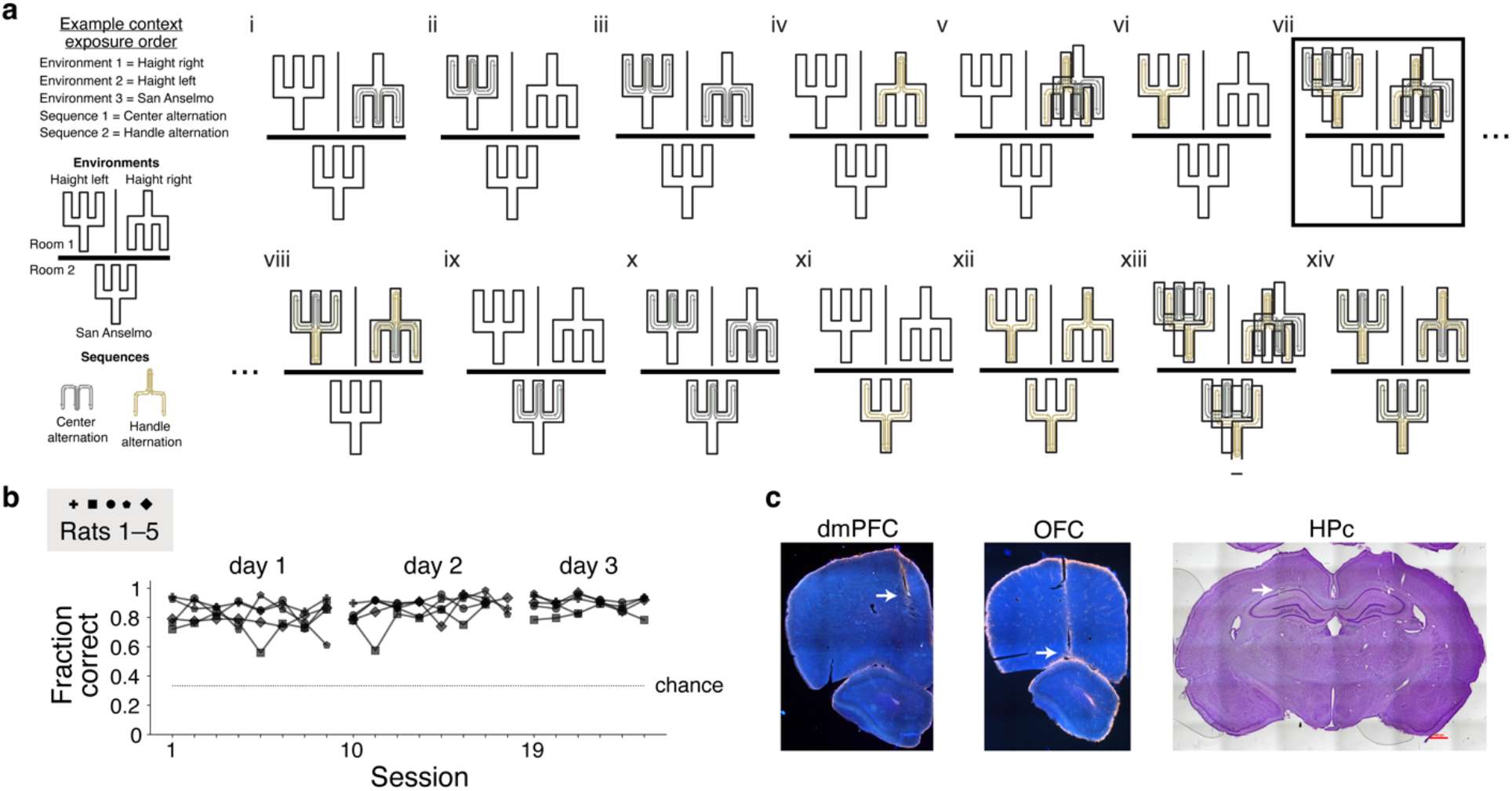
Behavioral training and histology. **a**, Timeline of exposures to the alternation task in different environments and with different sets of reward wells, illustrated for the case where environments are encountered in the order “Haight left”, “Haight right”, and “San Anselmo”, and sequences are encountered in the order “center alternation” (the designated home well is the center well) and “handle alternation” (the designated home well is the handle well). The black rectangle highlights the phase of behavior analyzed in this report; during this phase rats performed the task across two environments and sequences after having reached an 80% correct criterion in each of these contexts. **b**, Fraction of trials performed correctly in each session analyzed. Markers correspond to rats and lines connect sessions in a day. **c**, Example dmPFC, OFC, and HPc histological sections from one rat. Arrows point to electrode tracts.

**Extended Data Fig. 2.**
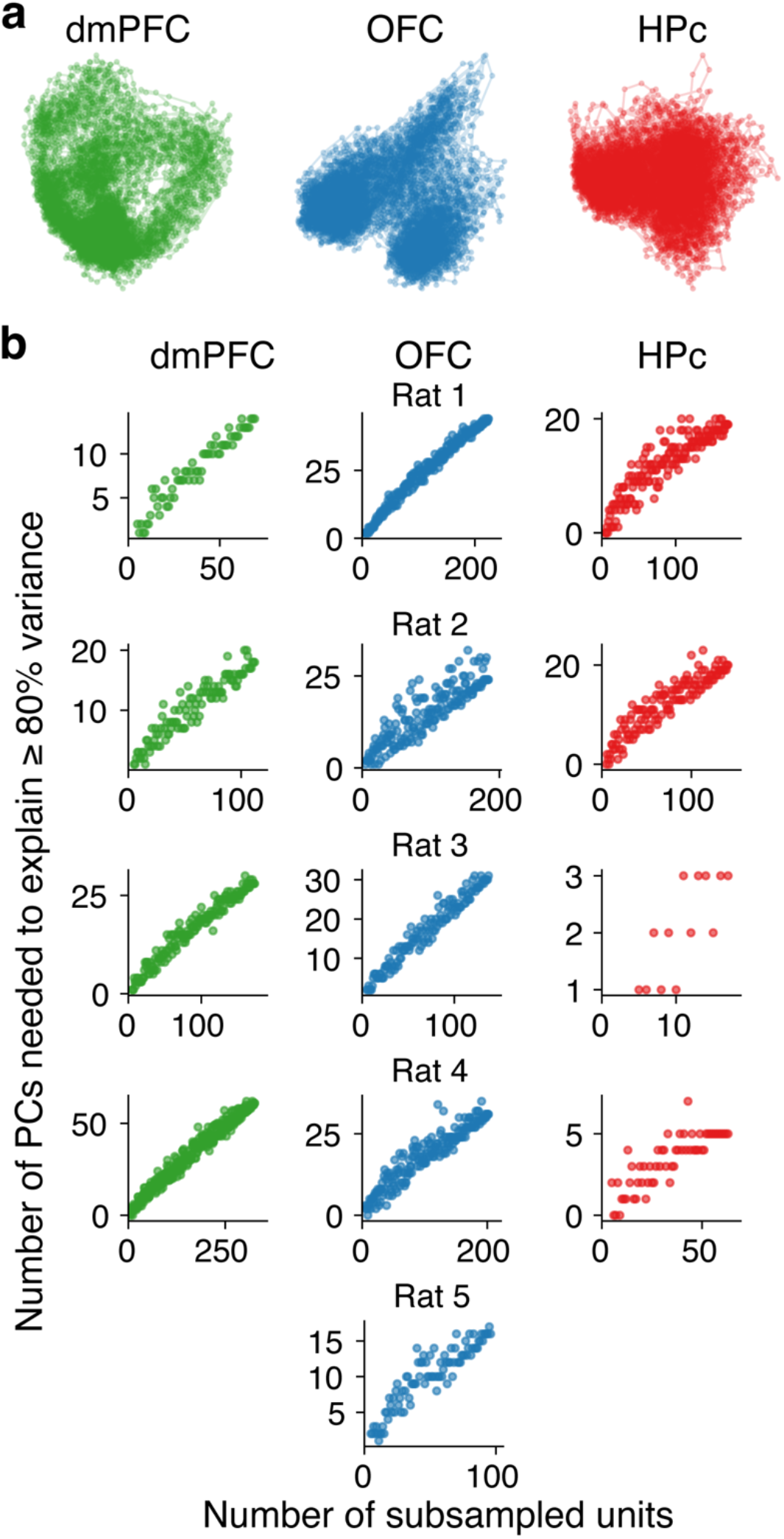
Population activity is not confined to a low-dimensional linear subspace. **a**, First two principal components (PCs) of a principal component analysis (PCA) on 100-ms firing rate vectors from simultaneously recorded single units in dmPFC (left), OFC (middle), and HPc (right) from an example behavioral session from one rat. **b**, Fraction of PCs needed to explain at least 80% of the variance in firing rate vectors in a brain region, as a function of subsampled ensemble size ranging from 5 to the number of simultaneously recorded units, for a representative session from each rat.

**Extended Data Fig. 3.**
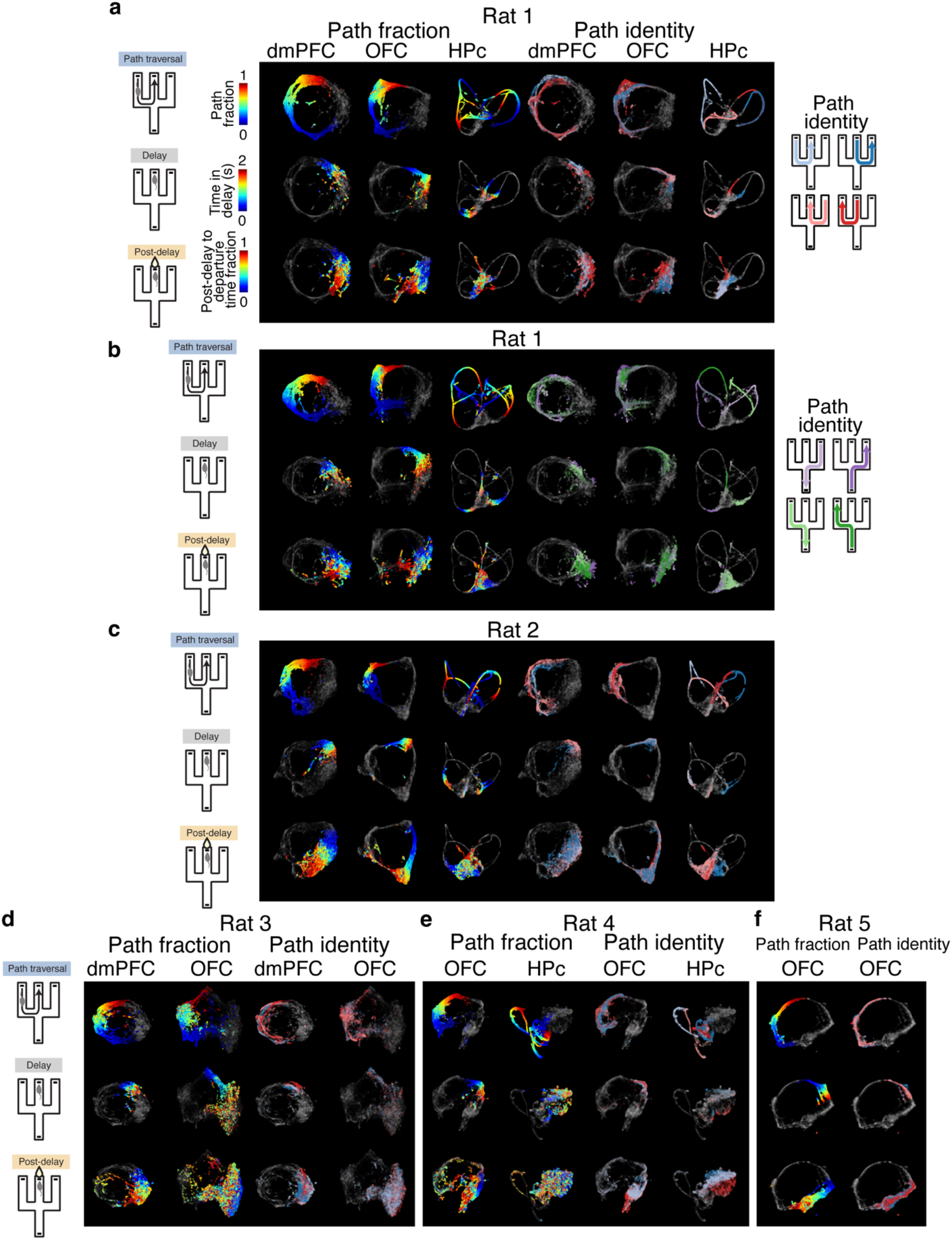
Population activity embeddings in different environments and with different sets of reward wells. **a**, UMAP embeddings of dmPFC (left), OFC (middle), and HPc (right) 100-ms firing rate vectors from the same rat shown in Fig. 2 but during a session in a different environment. Embedded firing rate vectors are colored according to the fractional distance along paths during the path traversal period (top left), time in the delay period (middle left), relative time in the post-delay period (bottom left), path identity during the path traversal period (top right), and the identity of the previously traversed path during the delay period (middle right) and post-delay period (bottom right). **b**, UMAP embeddings of dmPFC, OFC, and HPc firing rate vectors from the same rat shown in Fig. 2 during a session with a different set of reward wells. **c–f**, As in **a**, for rats 2–5.

**Extended Data Fig. 4.**
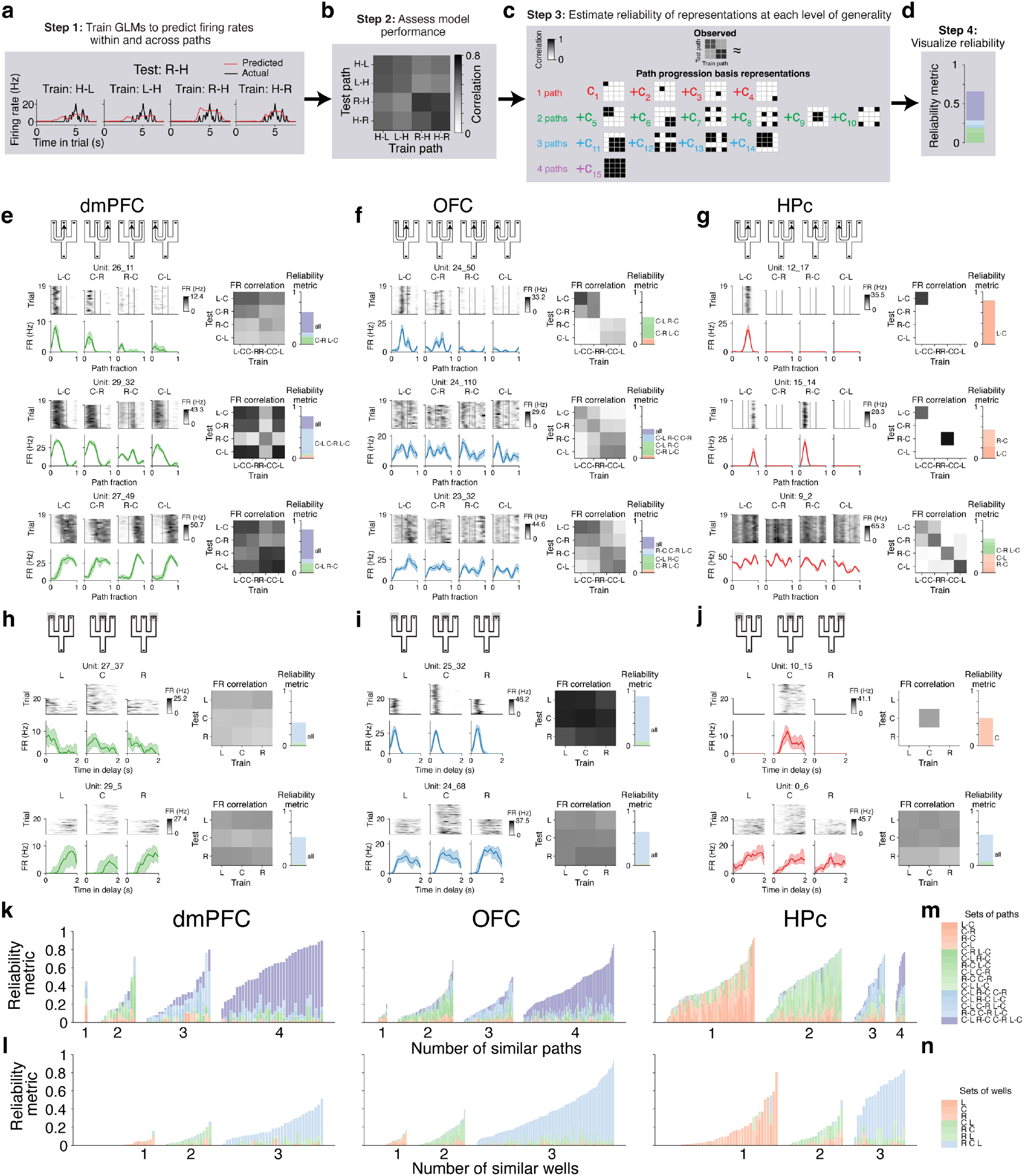
Quantifying the reliability of single unit representations of task progression at multiple levels of generality. **a–d,** Approach illustrated for path traversals using one single unit from a session with the handle alternation. **a,** Firing rates during a single trial on the path from the right well to the handle well (black), and firing rates predicted by a GLM (red) trained on each of the four potentially rewarded paths (first and second letters denote start and end wells; L = left well, R = right well, H = handle well). **b,** Matrix with Pearson correlation coefficient of actual and predicted firing rates for each pair of train and test path. **c,** Depiction of estimation of reliability of path progression representations at different levels of generality. The observed matrix of correlation coefficients in Step 2 is reconstructed as a positive linear combination of fifteen path progression basis vectors across four levels of generality. Each basis vector is the concatenation of entries in the expected matrix of correlation coefficients of actual and predicted firing rates for pairs of train and test paths given firing rate patterns that are perfectly reliable (invariant across trials) along one path (top row, “1 path”) or a specific set of two paths (second row, “2 paths”), three paths (third row, “3 paths”), or four paths (fourth row, “4 paths”), and unreliable along the remaining paths. c_1_–c_15_: positive weights that minimize the sum of squared differences between values in the observed and predicted correlation matrices. **d,** The magnitude of weights for each basis vector scaled by the number of paths to which it applies and normalized by the total number of paths with at least one spike (“reliability metric”) is shown as the height of colored bars. Reliability metrics are stacked, and those at higher levels of generality appear higher in the stack. Shades correspond to sets of paths as in **m**. **e–j,** In each panel: left: single trial firing rates (top) and trial average firing rates (with interquartile range; bottom) for the same units in Fig. 3a–f. Middle: Pearson correlation coefficients between firing rates along each path (matrix rows), and those predicted by a GLM trained on each of the four potentially rewarded paths (matrix columns). Right: stacked reliability metrics for basis vectors. As in **d**, height of bars corresponds to the magnitude of reliability metrics. Shades and abbreviations (shown for reliability metrics ≥ 0.1) are as in **m,n** and indicate the set of paths (**e–g** as in **m**) or wells (**h–j** as in **n**) represented as similar. **k–l,** Reliability metrics during the path traversal (**k**) or delay (**l**) period for all simultaneously recorded dmPFC (left), OFC (middle) and HPc (right) single units during the same example session shown in **e–j**. Each bar shows the stacked reliability metrics for one single unit as in **d**. Colors correspond to basis vectors as in **m,n**. Single units in each brain region are grouped by the level of generality for which the sum of reliability metrics at that level is greatest. Within groups, units are sorted in ascending order by the sum of all reliability metrics (i.e. bar height). **m,** Each shaded bar corresponds to one path progression basis vector. The set of paths along which perfectly reliable path progression representations are invariant is abbreviated to the right. Letters denote start and end wells of paths (L = left, R = right, C = center, H = handle). **m,** Each shaded bar corresponds to one delay period progression basis vector. The set of wells along which perfectly reliable delay period progression representations are invariant is abbreviated to the right. Letters denote wells (L = left, R = right, C = center, H = handle).

**Extended Data Fig. 5.**
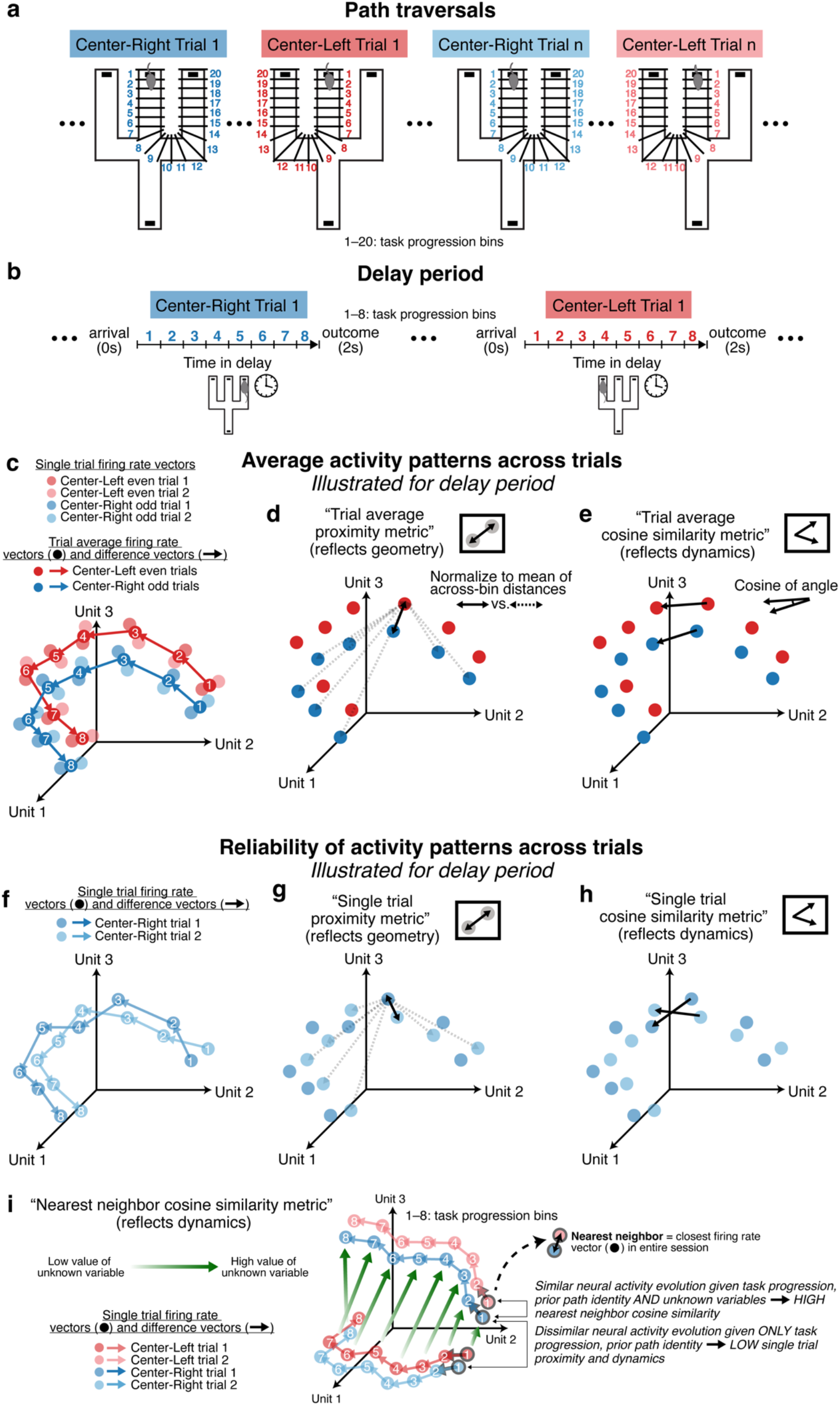
Quantifying encoding and reliability in population activity. **a**, Schematization of spatial task progression bins during path traversals. **b,** Schematization of temporal task progression bins during the delay period. **c–d**, Schematization of estimation of average population activity states and evolution during task progression. **c**, Single-trial (light shade) and even or odd trial-average (dark shade) firing rate vectors (dots) and difference vectors (arrows) in numbered task progression bins along the Center-Left (red) and Center-Right (blue) paths. **d**, Schematization of computing the “trial average proximity metric” in each task progression bin across the Center-Left and Center-Right paths, reflecting the extent to which average firing rate vectors tend to be closer together at a given task progression bin across paths in comparison to at different task progression bins across paths. **e**, Schematization of “trial average cosine similarity metric” applied to the Center-Left and Center-Right paths, reflecting the extent to which average firing rate vectors tend to evolve in similar directions at the same task progression bin across paths. **f**, Schematization of single-trial firing rate vectors (dots) and firing rate difference vectors (arrows) for two different trials (shades of blue) along the Center-Right path. **g**, Schematization of the “single trial proximity metric” applied to the Center-Right path, reflecting the extent to which firing rate vectors are reliably closer together at the same versus different task progression bins on different trials along the path. **h**, Schematization of the “single trial cosine similarity metric” applied to the Center-Right path, reflecting the extent to which firing rate vectors evolve in similar directions at a task progression bin on different trials along the path. **i,** Schematization of the “nearest neighbor cosine similarity” metric. A theoretical case where firing rate vectors evolve reliably as a function of task progression and previous path identity, but where evolutions are distinct given other hidden variables, is illustrated. Along different paths (Center-Left in red, and Center-Right in blue), firing rate vectors evolve differently for low or high values of an unknown variable, leading to low single trial proximity and dynamics along each path. However, firing rate vectors evolve similarly when values of the unknown variable are similar, and this is reflected in high cosine similarity of difference vectors of nearby states (“nearest neighbor cosine similarity”).

**Extended data Fig. 6.**
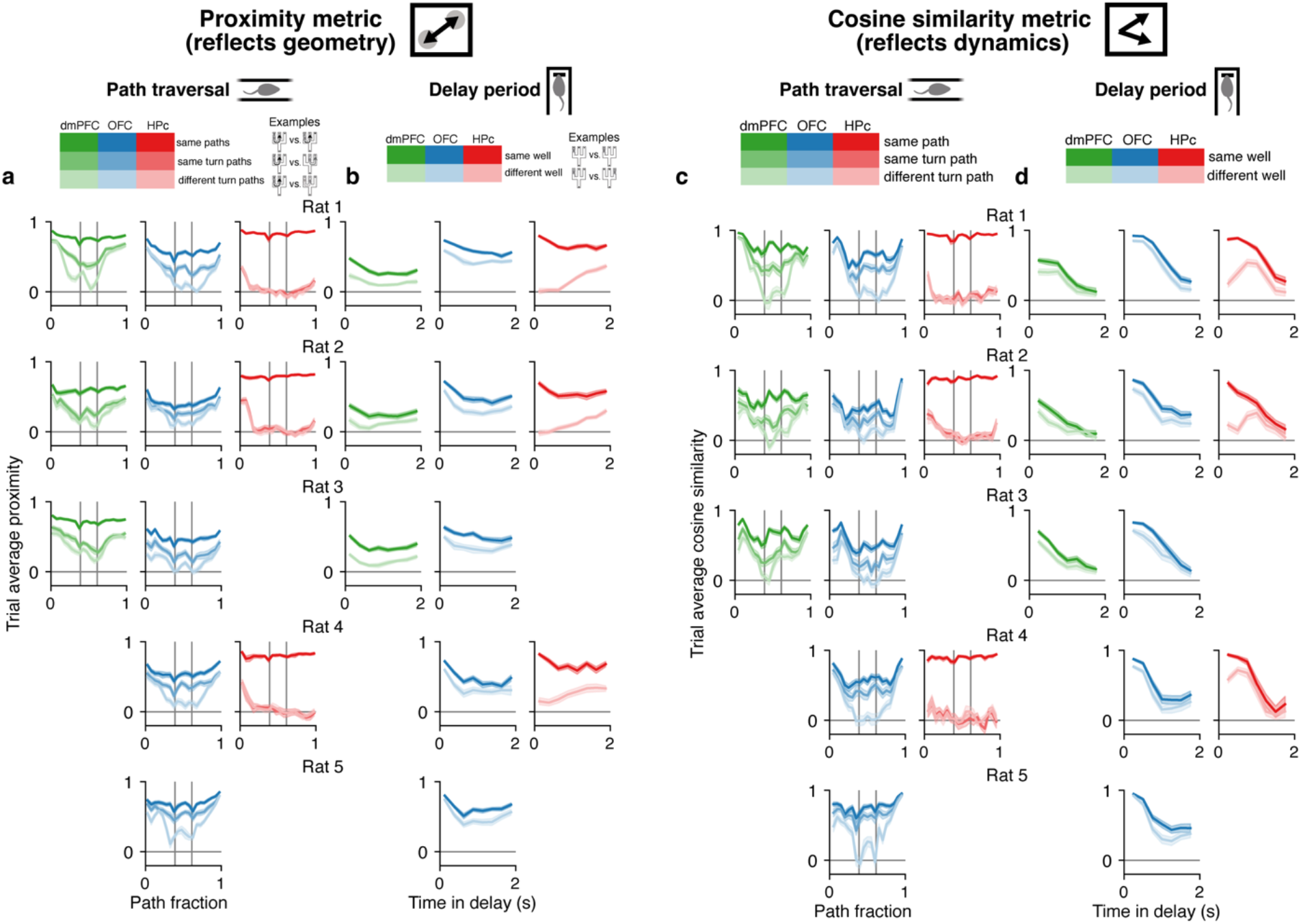
Average firing rate vector analysis for individual rats. **a**, As in Fig. 4a, shown for all rats. **b**, As in Fig. 4b, shown for all rats. **c**, As in Fig. 4e, shown for all rats. **d**, As in Fig. 4f, shown for all rats. In **a–d**, *n* = 22, 21, 21, 19, 22 sessions from rats 1–5. These plots illustrate that similar patterns of results are seen across animals.

**Extended Data Fig. 7.**
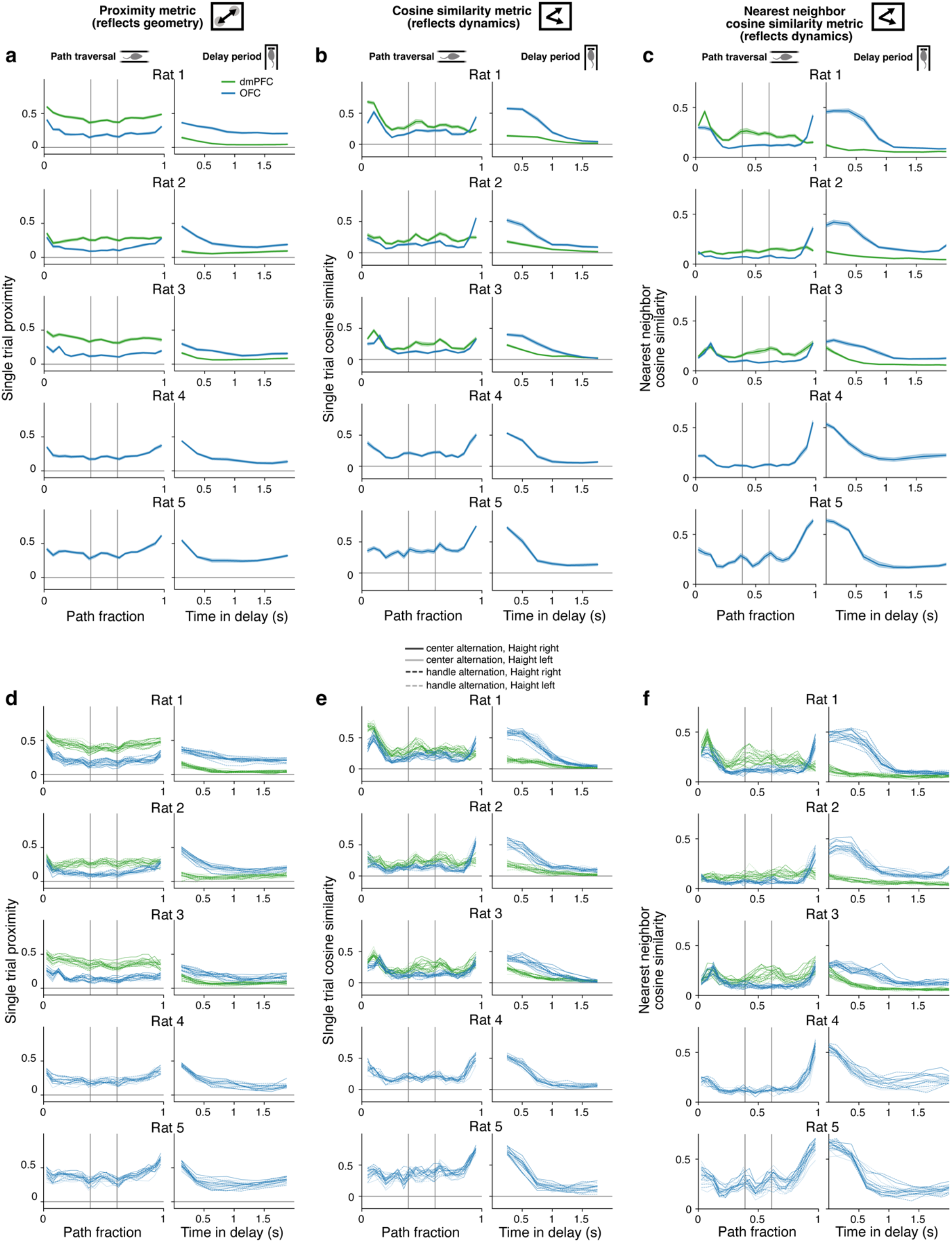
Single trial firing rate vector proximity and cosine similarity for individual rats. **a**, As in Fig. 4i, for all rats. **b**, As in Fig. 4k, for all rats. **c**, Cosine similarity of 100-ms firing rate difference vectors to their nearest neighbors in firing rate space across the entire recording session, averaged within spatial bins along paths (left) and time bins during the delay period (right) (mean and 95% CI, *n* = 22, 21, 21, 19, 22 sessions from rats 1–5). **d–f**, as in **a–c** but shown for single epochs. These plots illustrate that across animals, and within single sessions in different environments and with different sets of rewarded wells, measures of firing rate vector proximity and dynamics are more reliable during the path traversal in dmPFC, versus during the delay period in OFC.

**Extended Data Fig. 8.**
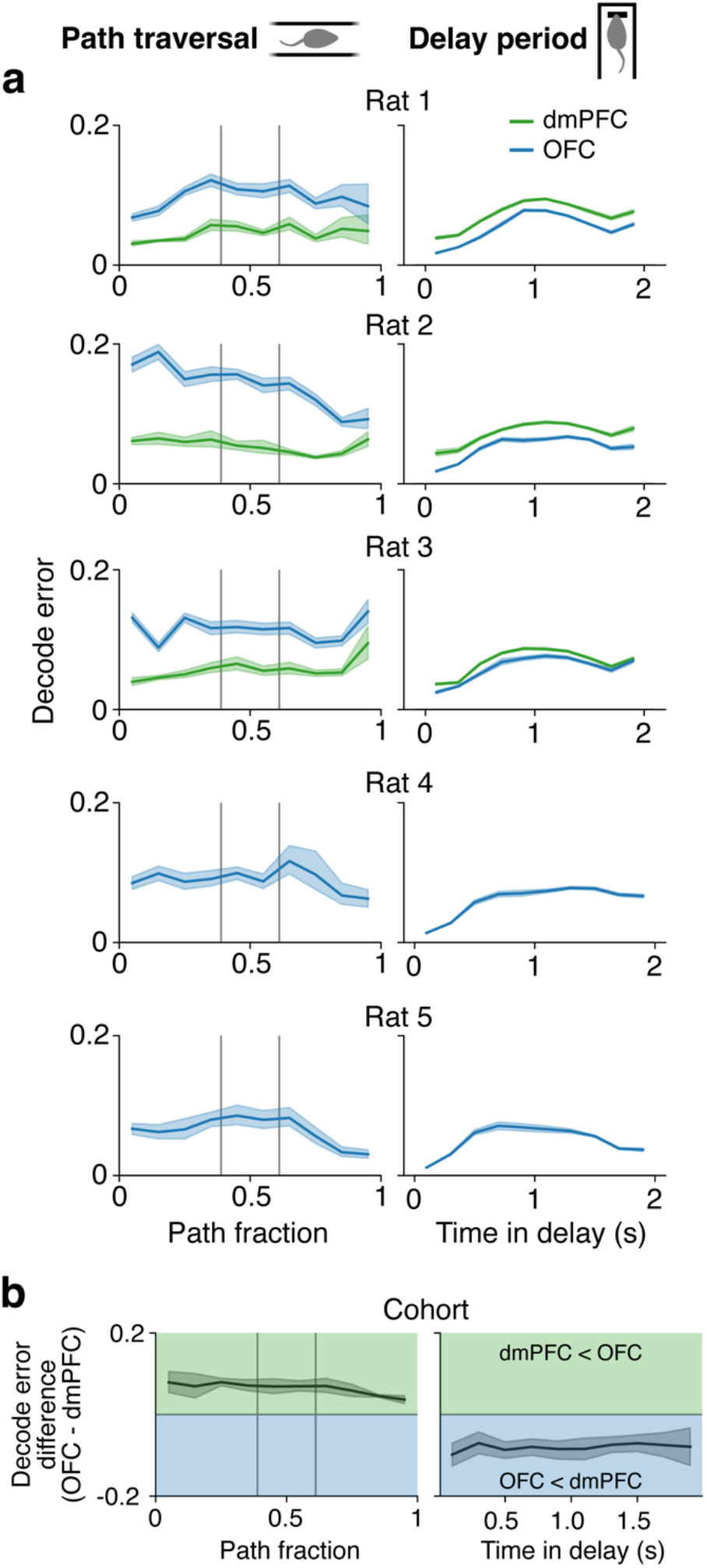
Task progression decoding is enhanced during path traversals using dmPFC ensembles and during the delay period using OFC ensembles. **a**, Absolute difference between actual path fraction (left) or time in the delay (right) and that predicted using a Bayesian decoder trained on random subsets of 50 units (mean and 95% confidence interval, *n* = 22, 21, 21, 19, 22 sessions from rats 1–5). Vertical lines mark maze junctions. **b**, Difference between decoding errors using OFC versus dmPFC 50-unit ensembles (mean and 95% CI, *n* = 64 sessions from *n* = 3 rats). Vertical lines mark maze junctions.

**Extended Data Fig. 9.**
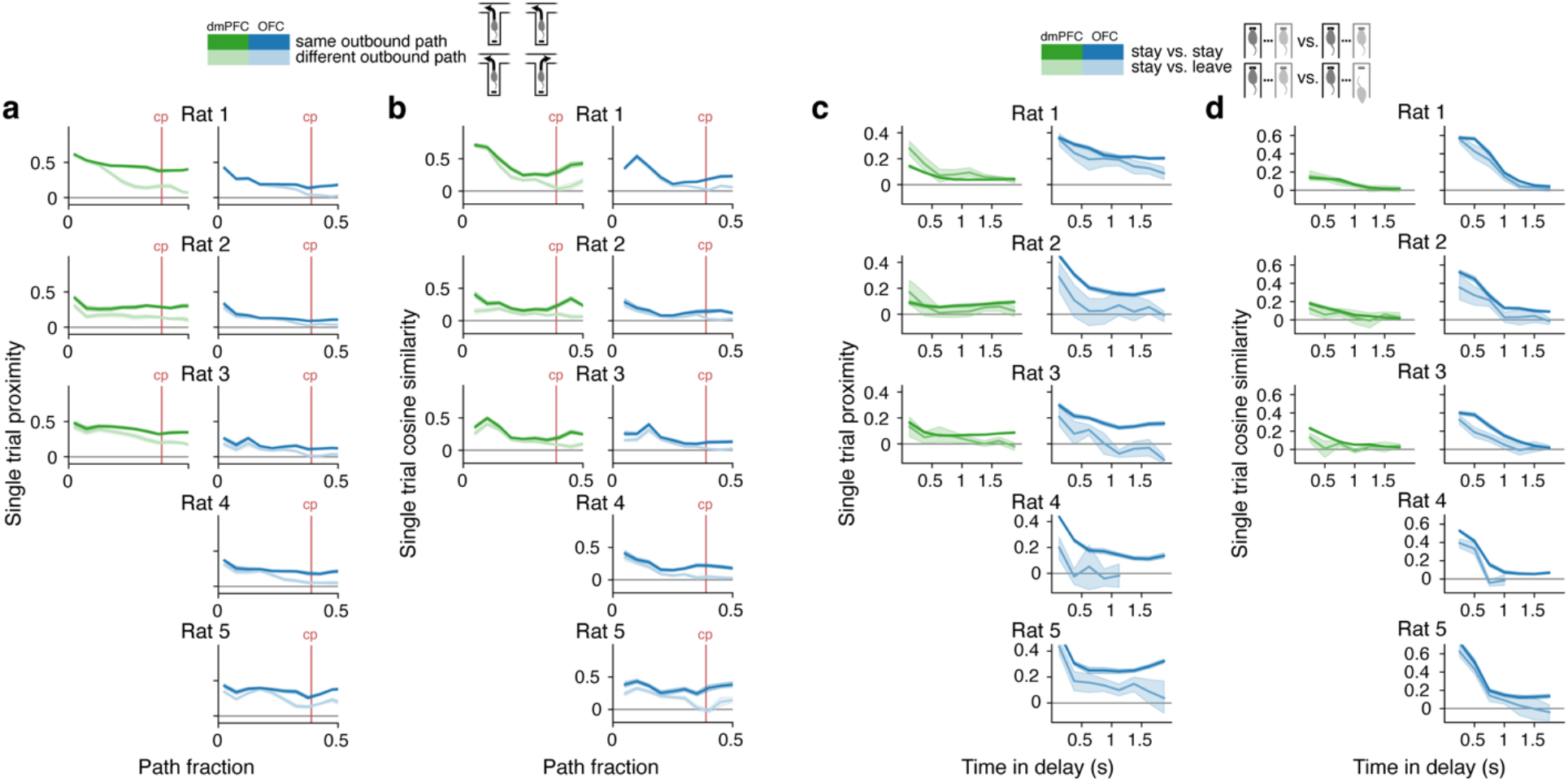
Regional enhancement in the reliable expression of firing rate states that distinguish upcoming actions in the dmPFC and waiting for versus forgoing future outcomes in the OFC. **a,** As in Fig. 5a, shown for all rats. **b,** As in Fig. 5d, shown for all rats. **c,** As in Fig. 5g, shown for all rats. **d,** As in Fig. 5j, shown for all rats. These plots show that single trial firing rate vector proximity consistently differs in association with future turn choice during the path period to a greater extent in dmPFC than in OFC. Conversely, single trial firing rate vector proximity consistently differs in association with the future decision to stay or leave during the delay period in OFC, but not in dmPFC. A similar trend is seen with single trial firing rate difference vector cosine similarity.

**Extended Data Table 1.** Mean number of single units (*± standard deviation*) recorded per session for each rat.

|  | dmPFC | OFC | HPc |
| --- | --- | --- | --- |
| **Rat 1** | 74 ± 9 | 224 ± 10 | 164 ± 11 |
| **Rat 2** | 111 ± 2 | 194 ± 9 | 160 ± 10 |
| **Rat 3** | 159 ± 19 | 130 ± 8 | 28 ± 9 |
| **Rat 4** | — | 203 ± 14 | 65 ± 12 |
| **Rat 5** | — | 103 ± 8 | — |

## REFERENCES

1. Cummings, J. L. Frontal-Subcortical Circuits and Human Behavior. Archives of Neurology 50, 873–880 (1993).

2. Buckley, M. J. et al. Dissociable Components of Rule-Guided Behavior Depend on Distinct Medial and Prefrontal Regions. Science 325, 52–58 (2009).

3. Rudebeck, P. H., Walton, M. E., Smyth, A. N., Bannerman, D. M. & Rushworth, M. F. S. Separate neural pathways process different decision costs. Nat Neurosci 9, 1161–1168 (2006).

4. Steinmetz, N. A., Zatka-Haas, P., Carandini, M. & Harris, K. D. Distributed coding of choice, action and engagement across the mouse brain. Nature 576, 266–273 (2019).

5. Stringer, C. et al. Spontaneous behaviors drive multidimensional, brainwide activity. Science 364, eaav7893 (2019).

6. Allen, W. E. et al. Thirst regulates motivated behavior through modulation of brainwide neural population dynamics. Science 364, eaav3932 (2019).

7. Miller, K. J., Botvinick, M. M. & Brody, C. D. Value representations in the rodent orbitofrontal cortex drive learning, not choice. eLife 11, e64575 (2022).

8. Rudebeck, P. H. et al. Frontal Cortex Subregions Play Distinct Roles in Choices between Actions and Stimuli. Journal of Neuroscience 28, 13775–13785 (2008).

9. Lak, A. et al. Orbitofrontal Cortex Is Required for Optimal Waiting Based on Decision Confidence. Neuron 84, 190–201 (2014).

10. Walton, M. E., Behrens, T. E. J., Buckley, M. J., Rudebeck, P. H. & Rushworth, M. F. S. Separable Learning Systems in the Macaque Brain and the Role of Orbitofrontal Cortex in Contingent Learning. Neuron 65, 927–939 (2010).

11. Jones, J. L. et al. Orbitofrontal Cortex Supports Behavior and Learning Using Inferred But Not Cached Values. Science 338, 953–956 (2012).

12. Costa, K. M. et al. The role of the lateral orbitofrontal cortex in creating cognitive maps. Nat Neurosci 26, 107–115 (2023).

13. Kennerley, S. W., Dahmubed, A. F., Lara, A. H. & Wallis, J. D. Neurons in the frontal lobe encode the value of multiple decision variables. J Cogn Neurosci 21, 1162–1178 (2009).

14. Sul, J. H., Kim, H., Huh, N., Lee, D. & Jung, M. W. Distinct Roles of Rodent Orbitofrontal and Medial Prefrontal Cortex in Decision Making. Neuron 66, 449–460 (2010).

15. Kennerley, S. W. & Wallis, J. D. Evaluating choices by single neurons in the frontal lobe: outcome value encoded across multiple decision variables. Eur J Neurosci 29, 2061–2073 (2009).

16. Feierstein, C. E., Quirk, M. C., Uchida, N., Sosulski, D. L. & Mainen, Z. F. Representation of Spatial Goals in Rat Orbitofrontal Cortex. Neuron 51, 495–507 (2006).

17. Samborska, V., Butler, J. L., Walton, M. E., Behrens, T. E. J. & Akam, T. Complementary task representations in hippocampus and prefrontal cortex for generalizing the structure of problems. Nat Neurosci 25, 1314–1326 (2022).

18. Zhou, J. et al. Evolving schema representations in orbitofrontal ensembles during learning. Nature 590, 606–611 (2021).

19. Zhou, J. et al. Rat Orbitofrontal Ensemble Activity Contains Multiplexed but Dissociable Representations of Value and Task Structure in an Odor Sequence Task. Current Biology 29, 897–907.e3 (2019).

20. Hirokawa, J., Vaughan, A., Masset, P., Ott, T. & Kepecs, A. Frontal cortex neuron types categorically encode single decision variables. Nature 576, 446–451 (2019).

21. Maharjan, D. M., Dai, Y. Y., Glantz, E. H. & Jadhav, S. P. Disruption of dorsal hippocampal – prefrontal interactions using chemogenetic inactivation impairs spatial learning. Neurobiology of Learning and Memory 155, 351–360 (2018).

22. Yu, J. Y., Liu, D. F., Loback, A., Grossrubatscher, I. & Frank, L. M. Specific hippocampal representations are linked to generalized cortical representations in memory. Nat Commun 9, 2209 (2018).

23. Kay, K. et al. A hippocampal network for spatial coding during immobility and sleep. Nature 531, 185–190 (2016).

24. Kennerley, S. W., Behrens, T. E. J. & Wallis, J. D. Double dissociation of value computations in orbitofrontal and anterior cingulate neurons. Nat Neurosci 14, 1581–1589 (2011).

25. Kato, S. et al. Global Brain Dynamics Embed the Motor Command Sequence of Caenorhabditis elegans. Cell 163, 656–669 (2015).

26. Churchland, M. M. et al. Neural population dynamics during reaching. Nature 487, 51–56 (2012).

27. McInnes, L., Healy, J. & Melville, J. UMAP: Uniform Manifold Approximation and Projection for Dimension Reduction. Preprint at http://arxiv.org/abs/1802.03426 (2020).

28. McNaughton, B. L., Barnes, C. A. & O’Keefe, J. The contributions of position, direction, and velocity to single unit activity in the hippocampus of freely-moving rats. Exp Brain Res 52, (1983).

29. Zhou, J. et al. Complementary Task Structure Representations in Hippocampus and Orbitofrontal Cortex during an Odor Sequence Task. Current Biology 29, 3402–3409.e3 (2019).

30. Yu, J. Y., Liu, D. F., Loback, A., Grossrubatscher, I. & Frank, L. M. Specific hippocampal representations are linked to generalized cortical representations in memory. Nature Communications 9, 2209 (2018).

31. Riceberg, J. S., Srinivasan, A., Guise, K. G. & Shapiro, M. L. Hippocampal signals modify orbitofrontal representations to learn new paths. Current Biology 32, 3407–3413.e6 (2022).

32. Tang, W., Shin, J. D. & Jadhav, S. P. Geometric transformation of cognitive maps for generalization across hippocampal-prefrontal circuits. Cell Reports 42, 112246 (2023).

33. Basu, R. et al. The orbitofrontal cortex maps future navigational goals. Nature 599, 449–452 (2021).

34. Baeg, E. H. et al. Dynamics of Population Code for Working Memory in the Prefrontal Cortex. Neuron 40, 177–188 (2003).

35. Sigala, N., Kusunoki, M., Nimmo-Smith, I., Gaffan, D. & Duncan, J. Hierarchical coding for sequential task events in the monkey prefrontal cortex. Proc. Natl. Acad. Sci. U.S.A. 105, 11969–11974 (2008).

36. Rubin, A. et al. Revealing neural correlates of behavior without behavioral measurements. Nat Commun 10, 4745 (2019).

37. Wilson, R. C., Takahashi, Y. K., Schoenbaum, G. & Niv, Y. Orbitofrontal cortex as a cognitive map of task space. Neuron 81, 267–279 (2014).

38. Musall, S., Kaufman, M. T., Juavinett, A. L., Gluf, S. & Churchland, A. K. Single-trial neural dynamics are dominated by richly varied movements. Nat Neurosci 22, 1677–1686 (2019).

39. Roumis, D. https://figshare.com/articles/software/physiology/4747438/2.

40. Chung, J. E. et al. Chronic Implantation of Multiple Flexible Polymer Electrode Arrays. Journal of visualized experiments: JoVE 152, (2019).

41. Sullivan, D. et al. Relationships between Hippocampal Sharp Waves, Ripples, and Fast Gamma Oscillation: Influence of Dentate and Entorhinal Cortical Activity. Journal of Neuroscience 31, 8605–8616 (2011).

42. Frank, L. M., Brown, E. N. & Wilson, M. Trajectory Encoding in the Hippocampus and Entorhinal Cortex. Neuron 27, 169–178 (2000).

43. Paxinos, G. & Watson, C. The Rat Brain in Stereotaxic Coordinates. (Elsevier, London, 2007).

44. Yatsenko, D., et al. DataJoint: Managing Big Scientific Data Using MATLAB or Python. http://biorxiv.org/lookup/doi/10.1101/031658 (2015) doi:10.1101/031658.

45. Buccino, A. P. et al. SpikeInterface, a unified framework for spike sorting. eLife 9, e61834 (2020).

46. Chung, J. E. et al. A Fully Automated Approach to Spike Sorting. Neuron 95, 1381–1394.e6 (2017).

47. Battaglia, F. P., Sutherland, G. R. & McNaughton, B. L. Local sensory cues and place cell directionality: additional evidence of prospective coding in the hippocampus. J Neurosci 24, 4541–4550 (2004).

48. Saravanan, V., Berman, G. J. & Sober, S. J. Application of the hierarchical bootstrap to multi-level data in neuroscience. Neuron Behav Data Anal Theory 3, (2020).

49. Truccolo, W., Eden, U. T., Fellows, M. R., Donoghue, J. P. & Brown, E. N. A Point Process Framework for Relating Neural Spiking Activity to Spiking History, Neural Ensemble, and Extrinsic Covariate Effects. Journal of Neurophysiology 93, 1074–1089 (2005).

50. Pillow, J. W. et al. Spatio-temporal correlations and visual signalling in a complete neuronal population. Nature 454, 995–999 (2008).

51. Park, I. M., Meister, M. L. R., Huk, A. C. & Pillow, J. W. Encoding and decoding in parietal cortex during sensorimotor decision-making. Nat Neurosci 17, 1395–1403 (2014).

52. McCullagh, P. & Nelder, J. A. Generalized Linear Models. (Routledge, 2019). doi:10.1201/9780203753736.

53. Seabold, S. & Perktold, J. Statsmodels: Econometric and Statistical Modeling with Python. Proceedings of the 9th Python in Science Conference 2010, (2010).

54. 4. The BFGS Method. in Iterative Methods for Optimization 71–86 doi:10.1137/1.9781611970920.ch4.

55. Chaudhuri, R., Gerçek, B., Pandey, B., Peyrache, A. & Fiete, I. The intrinsic attractor manifold and population dynamics of a canonical cognitive circuit across waking and sleep. Nat Neurosci 22, 1512–1520 (2019).

56. Koay, S. A., Charles, A. S., Thiberge, S. Y., Brody, C. D. & Tank, D. W. Sequential and efficient neural-population coding of complex task information. Neuron 110, 328–349.e11 (2022).

57. Denovellis, E. L. et al. Hippocampal replay of experience at real-world speeds. eLife 10, e64505 (2021).

